# Cross-fitted instrument: a blueprint for one-sample Mendelian Randomization

**DOI:** 10.1101/2021.05.05.441737

**Authors:** William R.P. Denault, Jon Bohlin, Christian M. Page, Stephen Burgess, Astanand Jugessur

## Abstract

Bias from weak instruments may undermine the ability to estimate causal effects in instrumental variable regression (IVR). We present here a simple solution for handling weak instrument bias by introducing a new type of instrumental variable called ‘cross-fitted instrument’ (CFI). CFI splits the data at random and estimates the impact of the instrument on the exposure in each partition. The estimates are then used to perform an IVR on each partition. We adapt CFI to Mendelian randomization (MR) and term this adaptation ‘Cross-Fitting for Mendelian Randomization’ (CFMR). A major advantage of CFMR is its use of all the available data to select genetic instruments, as opposed to traditional two-sample MR where a large part of the data is only used for instrument selection. Consequently, CFMR has the potential to enhance the power of MR in a meta-analysis setting by enabling an unbiased one-sample MR to be performed in each cohort prior to meta-analyzing the results across all the cohorts. In a similar fashion, CFMR enables a cross-ethnic MR analysis by accounting for ethnic heterogeneity, which is particularly important in consortia-led meta-analyses where the participating cohorts might be of different ethnicities. To our knowledge, there are currently no MR approach that can account for such heterogeneity. Finally, CFMR enables the application of MR to exposures that are rare or difficult to measure, which would normally preclude their analysis in the regular two-sample MR setting.

**Key messages:**

- We develop a new method to enable an unbiased one-sample Mendelian Randomization.
- The new method provides the same power as the standard two-sample Mendelian Randomization approach and does not require summary statistics from a genome-wide association study in an independent cohort.
- Our approach enables a cross-ethnic instrumental variable regression to account for heterogeneity in a sample consisting of multiple ethnicities.

## 1. Introduction

There are many successful applications of two-sample MR for inferring causal relationships, but its use is limited when studying exposures not typically studied by large consortia. Performing a two-sample MR using an exposure with no previously reported findings from genome-wide association studies (GWASes) requires coordinating analysis between at least two large non-overlapping cohorts from a similar population, which is time-consuming and often unfeasible. Furthermore, the requirement for the two cohorts to stem from a similar population limits the application of two-sample MR to cohorts comprising different ethnicities that have not yet been adequately genotyped.

To address these shortcomings, we present here a solution that relies on a simple modification of the two-stage least square (2SLS) procedure (1) that satisfies the main assumptions of two-sample MR (the samples must be non-overlapping and must stem from a similar population) while using only a single dataset for instrument selection. This modification exploits the concept of cross-fitting (CF) from the debiased machine-learning (DML) approach recently proposed by Chernozhukov and colleagues (2). In essence, we construct a new type of instrument based on CF, which we termed ‘cross-fitted instrument’ (CFI), that allows a conservative estimation of the causal effect of an exposure on an outcome. Other ideas analogous to CF can be found in the earlier works by Angrist and colleagues (3; 4). CFI differs from these approaches by its use of a data-splitting procedure that allows all the available data to be used in instrument selection, thus eliminating the need for sub-sampling (3). As a result, CFI is able to reduce the computational burden compared to the jackknife procedure by Angrist *et al.* (4).

We call the adaption of CFI to MR ‘Cross-Fitting for Mendelian Randomization’ (CFMR) and show that it uses more data than traditional two-sample MR when estimating the causal effect of an exposure on an outcome. Other works on similar topics have recently appeared in the literature (5; 6; 7); however, some of those investigations are not yet peer reviewed (archived), are limited by the need for a large sample size (5), or are restricted to inverse-variance two-sample weighting MR (6; 7). By contrast, CFMR can be applied to small sample sizes and is easily adaptable to a polygenic risk score (PRS) setting. Finally, our work is also closely related to the recently proposed ‘causal gradient boosting’ approach by Bakhitov and Singh (8), which is also at the preprint stage. Bakhitov and Singh (8) also use CF in the context of non-linear instrumental variable regression involving a small number of instruments. CFMR differs from that approach in that it is restricted to linear instrumental variable regression and can handle a larger number of instruments.

## 2. Methods

In their pioneering work, Chernozhukov and colleagues (2) proposed two causal estimators, DML1 and DML2, that are asymptotically equivalent. We present here their MR counterparts, CFMR1 and CFMR2, that are also asymptotically equivalent. For extensive details regarding the optimal selection of genetic instruments, readers are referred to the work by Hemani and colleagues (9). To simplify further, we have only considered here the case where the genetic instrument does not exhibit pleiotropic effects (10). This is because CFMR can easily be adapted to recently developed MR methods that allow the use of pleiotropic genetic instruments, such as MR-PRESSO (11) and IVs based on penalized regression (12; 13; 14), which enable valid inference even when 50% of the SNPs are not valid instruments.

In the subsections below, we explain the concept of *K*-fold CFI and define the estimators CFMR1 and CFMR2. To ease comprehension, we also provide a simple example of the 2-fold CFMR in the Supplementary Material, in the subsection called ‘2-fold CFMR’.

### 2.1. CFMR

Let *Y* be a continuous outcome, *X* a continuous exposure, and *Z* a vector of size ≥ 1 containing the instruments. We assume that *Y*, *X* and *Z* are connected through the following linear regression models:

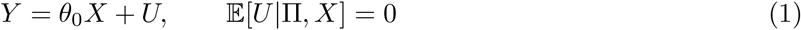

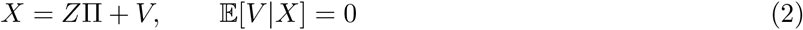

The parameter of interest is *θ*_0_ is the causal effect of X on Y, Π is the vector of regression coefficients for the instruments, *U* and *V* are two correlated error terms, and 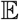 is the expectation operator.

#### 2.1.1. K-fold CFI

A CFI based on *K* ≥ 2 splits is referred to as a *K*-fold CFI, which can be described as follows. Let us consider *K*-fold random partitions of the observation indices [*N*] = (1, …, *N*), where the size of each fold is 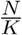. We refer to these partitions as 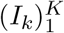. For each *k* ∈ (1, …, *K*), we define the complementary of the partition *I_k_* as 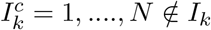. For each *k*, we select *n_k_* independent variants 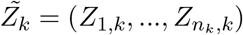 by performing a GWAS of *X* using the data in *I_k_*. In our application to a real dataset (see the subsection ‘Application of CFMR to a real dataset’ further below), 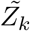 is the output of the clumped GWAS result of *X* using data with an index in *I_k_*.

We then use these *n_k_* variants to build *pred_k_* (a predictor of *X*) and use the data with an index in *I_k_* as a training set. The predictor *pred_k_* can be based on any machine learning/statistical method suitable for building instrumental variables, such as the least absolute shrinkage and selection operator (LASSO) (12) or PRS (15), but non-linear methods such as the generalized random forest (16) can also be applied. For each *k*, we define the CFI of *X* on 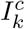 as:

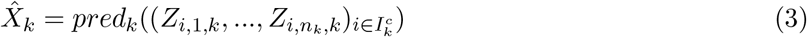

Where *Z_i,l,k_* is the variant *Z_l,k_* of individual *i*. A CFI on 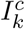 is the prediction of *X* on 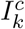 using a predictor of *X* trained using data with an index in *I_k_*. For 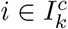, we denote the predicted exposure of individual *i* using *pred_k_* as 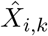. Finally, the *K*-fold CFI, 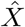, is defined as:

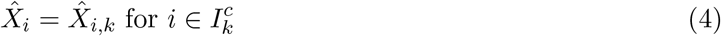

### 2.2. CFMR

The CFMR1 estimate of *θ*_0_ is defined as:

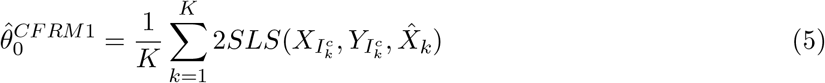

Where 2*SLS* is the 2SLS estimator (17):

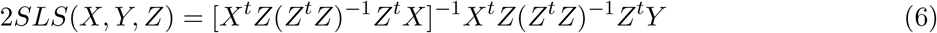

This estimate corresponds to step 4 in panel b in Figure 1. CFMR1 consists of performing an IVR on the complementary partition 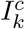 using 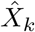 as instrument. We then average the estimates of these IVRs to obtain the final estimate.

**Fig. 1.**
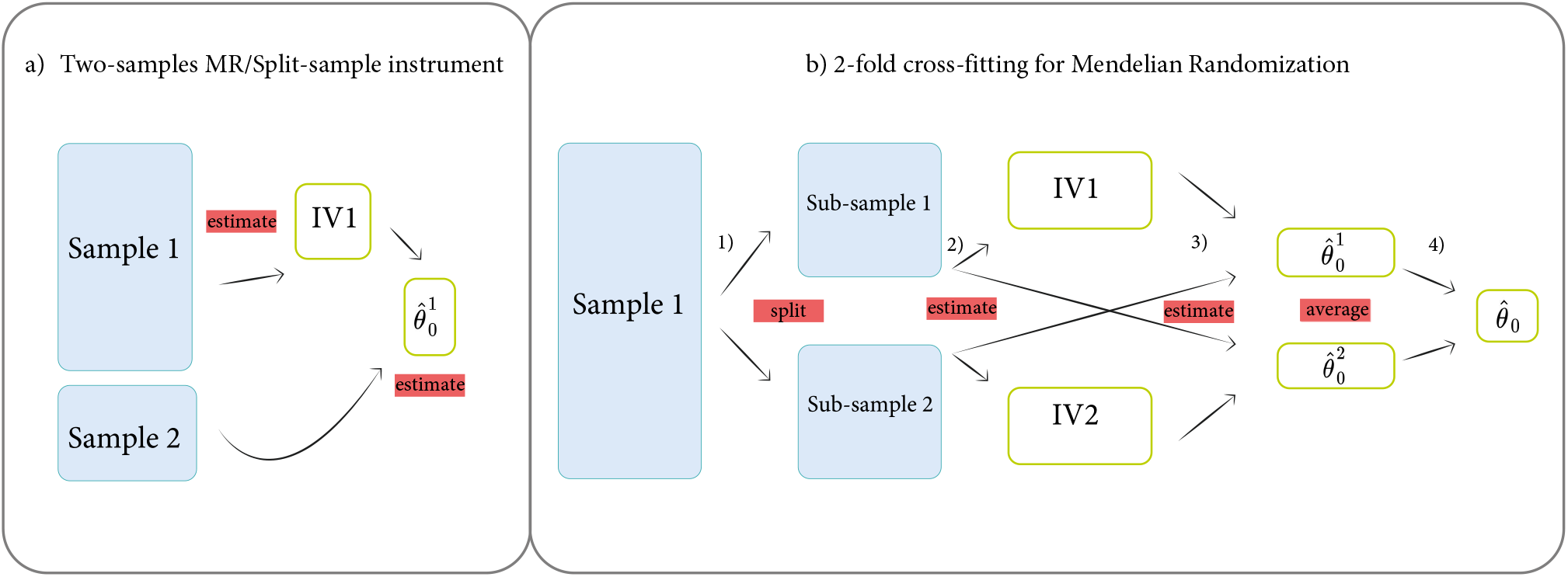
Schematic overview of two-sample MR and two-fold CFMR. Panel (a) shows the two-sample MR set-up in which the first sample is used to build the instrument while the second sample is used to estimate the causal effect. Panel (b) shows the two-fold CFMR set-up. Step 1 in panel b) describes the random splitting of the dataset. In step 2, two separate GWASes are performed: the first using sub-sample 1 and the exposure; the second using sub-sample 2 and the exposure. The predictors of the exposure are subsequently built based on sub-sample 1 (IV1) and sub-sample 2 (IV2). Step 3 refers to the 2SLS using IV1 on sub-sample 2 and IV2 on sub-sample 1. Finally, in step 4, the two 2SLS from step 3 are simply averaged.

The CFMR2 estimate of *θ*_0_ is simply defined as:

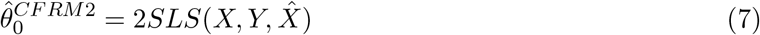

In essence, CFMR2 consists of performing a single IVR on the entire dataset using 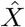 as instrument. In both CFMR1 and CFMR2, 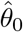 is asymptotically normally distributed around *θ*_0_ (see (3; 4)).

#### Finite sample unbiased estimation of 2SLS

As pointed out by Nagar (18), the expected bias of a 2SLS estimate is given by the expectation of 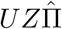, which is the following when using standard 2SLS:

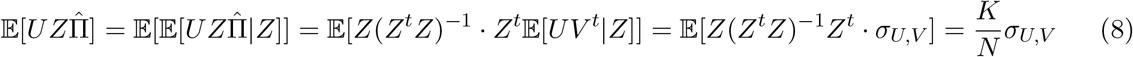

where 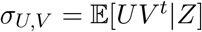 is the covariance of the error terms in the first and second-stage regression. Similar to the argument set forth by Angrist *et al.* (4), given that we use a CF procedure, 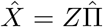 is by design independent of *U* and the error terms are independent. Hence, 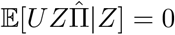.

The CFMR1 and CFMR2 estimates converge to their true value at a rate of 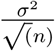, regardless of the strength of the instrument and the number of splits (4). The convergence speed of CFMR is the same as that of the standard two-sample MR (19), which implies that the two approaches have the same asymptotic power. We show via simulations (see Supplementary section 3.1) that CFMR and two-sample MR have equal power (Supplementary Figure S1 and Supplementary Table 1). To further assess the behavior of CFMR, in terms of its handling of bias, type I error and power, we performed a set of additional simulations and provide the results in the Supplementary Material. Specifically, we demonstrate that CFMR is conservative even under extreme scenarios in which one-sample MR is heavily biased (see Supplementary section 3.2).

## 3. Application of CFMR to a real dataset

We applied CFMR2 to a dataset comprising mother-child duos from the Norwegian Mother, father, and Child Cohort Study (MoBa) (20) to re-examine the well-established effect of maternal pre-pregnancy BMI on offspring’s birth weight (21). We chose CFMR2 over CFMR1 because, similarly to DML1 and DML2, CFMR2 exhibits a better finite sample size performance than CFMR1 (see (2)). After applying the quality control criteria outlined in the section ‘Study description’ in the Supplementary Material, 26, 896 complete mother-child duos with genotype and phenotype data remained for the current analyses. The maternal genotype was used to build the instrument for pre-pregnancy BMI. As additional criteria, we assumed random mating between parents and independence between mothers (i.e., no sibships) (22). CFMR was run on the 26, 896 mother-child duos using 10 random splits. Thus, 10 separate GWASes of pre-pregnancy BMI were performed, with each GWAS encompassing 24, 210 randomly selected mothers (24, 210 is about 90% of the original 26, 896 mothers). As our sample is relatively modest in size, we only used the first three principal components (PCs) to adjust for population stratification in each GWAS.

The Manhattan plots of the 10 GWASes are provided in Supplementary Figures 9–18. The results across the GWASes are similar and show a systematic replication of the top hits previously identified in two large GWAMAs of BMI (23; 24), which included SNPs in the genes *FTO*, *TMEM18*, and *MC4R* (8). Furthermore, we clumped the results of each GWAS and selected SNPs with a P-value below 10^−6^ to build a predictor of maternal pre-pregnancy BMI using LASSO. We then built the CFI by predicting each mother’s pre-pregnancy BMI using a predictor trained on a dataset that does not contain the data to be predicted. We repeated this procedure using different P-value thresholds ranging from 10^−3^ to 10^−8^ to build a CFI for each of these thresholds.

In addition, to compare CFMR with one-sample MR, we also performed a GWAS of pre-pregnancy BMI using the entire dataset of 26, 896 mother-child duos. Again, we clumped the results of the GWAS and selected SNPs with a P-value below 10^−6^ to build a predictor of maternal pre-pregnancy BMI by applying LASSO to the entire dataset. We then used the prediction of the predictor of maternal pre-pregnancy BMI as instrument. We refer to the estimation of the effect of maternal pre-pregnancy BMI on birth weight as one-sample MR estimation. Similar to the analyses above for the truncated dataset, we used different P-value thresholds ranging from 10^−3^ to 10^−8^ and estimated the effect of maternal pre-pregnancy BMI for each of these thresholds.

### 3.1. Application results

Except for the CFI constructed using SNPs with a P-value below 10^−8^, CFIs of maternal pre-pregnancy BMI explained about 1% of the variance in pre-pregnancy BMI (Table 1). When testing the CFIs for association with potential confounders, such as maternal age and pre-pregnancy maternal smoking (21), we found no evidence of an association between these variables and the CFIs constructed using SNPs with a P-value below 10^−6^. By contrast, for CFIs constructed using SNPs with a P-value threshold larger than 10^−6^, we observed moderate to strong associations with these variables. The results are summarized in Supplementary Table S6.

**Table 1:** CFMR estimates of maternal pre-pregnancy BMI on offspring birth weight per 1 SD increase in maternal pre-pregnancy BMI. The column “−*log*_10_ SNP P-value” corresponds to the cutoff used to build the CFI. The column “1SMR estimate” corresponds to the estimation using one-sample MR. The column “1SMR Std. error” corresponds to the standard error of the estimation based on one-sample MR. The column “Variance explained” corresponds to the pre-pregnancy variance explained by the CFI. The column “SNPs per fold” corresponds to the average number of SNPs with a P-value below a given threshold after clumping the output of each GWAS of maternal pre-pregnancy BMI. The column “Selected SNPs per fold” corresponds to the average number of SNPs selected by LASSO to build the - instrument in each fold.

| $-\log_{10}$ SNP<br>P-value | 1SMR<br>estimate | 1SMR<br>Std. error | CFI variance<br>explained (%) | CFMR<br>estimate | CFMR<br>Std. error | P-value | 95% CI | SNPs<br>per split |
| --- | --- | --- | --- | --- | --- | --- | --- | --- |
| -3 | 84.4 | 4.4 | 1.112 | 101.6 | 25.0 | 0.00005 | 52.6-150.6 | 1798 |
| -4 | 88.6 | 5.8 | 1.102 | 113.8 | 26.3 | 0.00002 | 62.2-165.5 | 624 |
| -5 | 88.1 | 8.4 | 1.101 | 94.3 | 26.6 | 0.00038 | 42.3-146.4 | 198 |
| -6 | 94.5 | 12.3 | 1.112 | 82.4 | 24.9 | 0.00093 | 33.6-131.2 | 52 |
| -7 | 85.6 | 16.3 | 0.951 | 73.4 | 27.0 | 0.00657 | 20.5-126.2 | 22 |
| -8 | 108.1 | 19.2 | 0.044 | 87.0 | 38.4 | 0.02351 | 11.7-162.3 | 6 |

For each CFI, we performed 2SLS to estimate the causal effect of maternal pre-pregnancy BMI on offspring birth weight. To follow the approach of Tyrrell and colleagues (21), we also adjusted for maternal age and fetal sex in each of these 2SLS. The CFMR estimates remained similar across the different CFIs (Table 1, Supplementary Figure S31, and Supplementary Table S5). Table 1 summarizes the CFMR estimates generated by using the CFI based on SNPs with a P-value below 10^−7^. We used this CFI because it showed no association with the potential confounders and explained a relatively large fraction of maternal pre-pregnancy BMI (0.91%).

Our CMR results indicated that a genetically-predicted increase of 1 SD of maternal pre-pregnancy 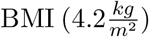 was associated with an increase in offspring birth weight of 73.35 g (95% CI: 20.46−126.24, *P* = 6.56 × 10^−6^), which corresponds to an increase in offspring birth weight of 17.42 g (95% CI: 4.86 − 29.98, *P* = 6.56 × 10^−6^) per unit increase of genetically-predicted maternal pre-pregnancy BMI. These CFMR estimates are similar to the observational associations in our data set of 81.90 g (95% CI: 75.93 − 87.88, *P* < 10^−16^) increase in offspring birth weight per 1 SD higher maternal pre-pregnancy BMI (19.44 g, 95% CI: 18.03 − 20.87, *P* < 10^−16^).

## 4. Discussion

This study presents a new type of IV, termed CFI, that is readily adaptable to an MR setting. The main advantage of CFMR over regular two-sample MR is its ability to perform two-sample MR using a single sample, which allows its application to considerably smaller sample sizes than is feasible by two-samples MR. Furthermore, CFMR ensures that the population assumptions of MR are fulfilled. Additional advantages of CFMR include affording the same power as two-sample MR and allowing estimates from multiple CFMRs to be meta-analyzed while taking into account heterogeneity between the different estimates (Figure 2). As CFMR is modular, it lends itself easily to parallel computing and can therefore be used in conjunction with many statistical methods to build multiple variant scores for downstream analyses, such as polygenic risk scores (25) or LASSO-based instruments (26; 12; 13; 14).

**Fig. 2.**
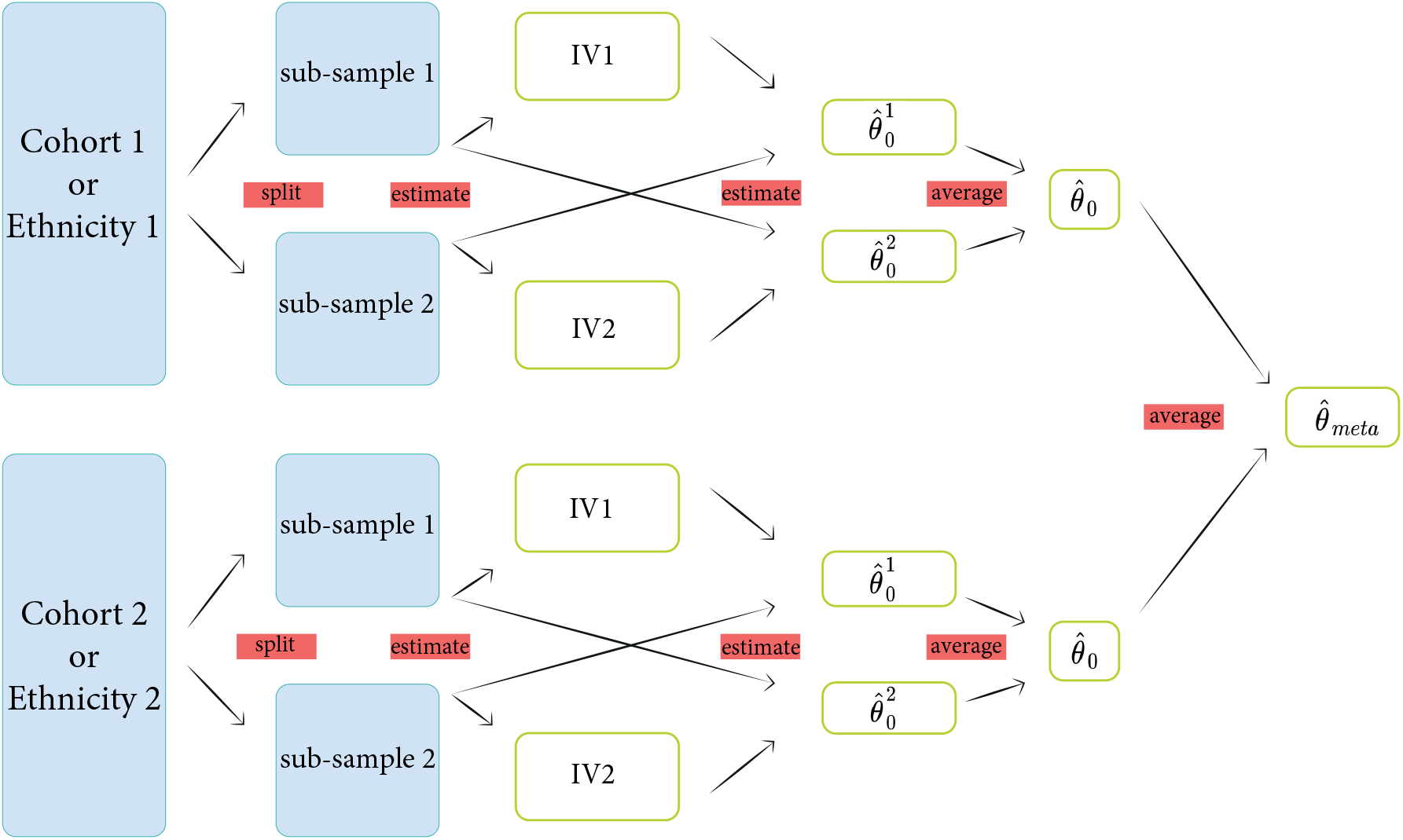
Application of CFMR to a sample composed of multiple ethnicities.

Compared to two-sample MR, in which one sample is used to build the instrument while the other is used to test for association, CFMR allows an unbiased estimation of causal effect using two samples from the same source population. If two separate samples are available for analysis, CFMR can be applied to each sample separately followed by a meta-analysis of the results. As meta-analyzing the results increases the active sample size of the study compared to a regular two-sample MR, CFMR can potentially enhance the power of regular MR analyses. By the same token, CFMR can easily handle multiple ethnicities in the same sample by enabling separate analyses of each ethnicity followed by a meta-analysis of the results. This type of cross-ethnic MR would enable accounting for potential heterogeneous effects of the exposure on the outcome across different sub-populations.

Chernozhukov and co-workers (2) reported that the number of splits has a negligible impact on the asymptotic convergence speed of the DLM methods. We observed the same trend in our data: the number of splits had no appreciable effect on the convergence speed of the exposure predictors in CFMR. However, as the power of 2SLS depends heavily on the prediction accuracy of the IV (19), it is critical to obtain a good genetic predictor of the exposure. Two to five splits may be sufficient if a large sample size (≥ 100, 000) is available for analysis and the exposure under study is highly heritable (e.g., if a few SNPs have large individual effects on the exposure). However, for more complex traits and smaller sample sizes, it may be beneficial to increase the number of splits to improve the predictive performance of the exposure predictors (2). As a rule of thumb, we recommend using CFMR with ten splits. When the exposure is particularly difficult to predict or the sample size is limited (≤ 5000), or both, using 20-30 splits may provide some improvement. Moreover, we recommend using CFMR2 in most practical settings, as also recommended by Chernozhukov and colleagues (2). The rationale for this is that CFMR1 is asymptotically equivalent to CFMR2, but as CFMR1 is an average of estimates based on small datasets, which can be noisy, CFMR1 tends to be less powerful than CFMR2 for small sample sizes. In our presentation of CFMR, we suggest clumping the GWAS results prior to building the IV. However, the application of other steps, such co-localization (27; 28) or whole-genome (LASSO) regression (29), may also be worth pursuing in this context.

Our simulations indicate that CFMR remains unbiased as long as the sample size is sufficiently large (≥ 100, 000). When the variance explained by the instruments is small (e.g., *h*^2^ ≤ 1%), CFMR is biased toward the null, which makes it a conservative approach for causal estimation. Another attractive feature of CFMR is its good control of type I error even when no instrument is associated with the exposure (i.e., *h*^2^ = 0). This tight error control is partially due to the standard errors of CFMR being too large for small sample sizes (≤ 10, 000). This is not unexpected, given that the standard error for 2SLS estimates is based on normal approximation, which may not be valid for small sample sizes (17). Obtaining narrower confidence intervals for weak CFIs may increase the power of CFMR for analyzing complex traits and exposures that have low heritability.

Future research should focus on deriving the CFMR distribution for weak instruments using robust variance estimates. Aside from challenges related to limited sample size, our simulations showed bias toward the null for weak instruments (*h*^2^ ≤ 1%), which is as expected given that we do not assume to know the causal variant *a priori*. For instance, when the total variance explained by the SNPs is 0.1%, each SNP explains only around 0.02% of the exposure heritability. Therefore, it becomes challenging for LASSO (26) to select causal variants among the 300 variants used in our simulations. Another limitation of CFMR is pleiotropy, which not only affects CFMR but also the vast majority of other MR-based methods, perhaps with the exception of CAUSE (10) and MR-PRESSO (11).

In our application of CFMR to real data, we estimated the effect of maternal pre-pregnancy BMI on offspring birth weight in the Norwegian MoBa study. Predictors of maternal pre-pregnancy BMI explained, on average, 11.4% of the variance in maternal pre-pregnancy BMI in the training sets (Supplementary Figures S20–S30). In comparison, our CFI based on SNPs with a P-value below 10^−7^ explained only 0.915% (P-value ≤ 10^−16^) of the variance. The difference in variance explained by the CFIs in the training versus test sets illustrates how CFMR can circumvent the problem of overfitted instruments (see also Supplementary Figures 19–30). The variance explained by our CFIs is smaller (between 0.915% and 1.112%) than the variance explained by the IV used by Tyrrell and colleagues (1.8%) (21). Interestingly, our estimates of maternal pre-pregnancy BMI on offspring birth weight are similar to those reported by Tyrrell *et al.* (21). Notably in the Tyrrell *et al.* study, 1 SD increase in maternal pre-pregnancy BMI (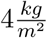 in Tyrrell *et al.* and 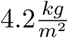 in our analysis) corresponded to an increase of 55 g (95% CI: 17 − 93) in offspring birth weight (73.35 g (95% CI: 20.46 − 126.246) in our analysis). In the Tyrrell *et al.* study, the observational association corresponded to 62 g (95% CI: 56 − 70) increase in offspring birth weight per 1 SD of higher maternal pre-pregnancy BMI.

Moreover, the size of the Tyrrell *et al.* study is very similar to ours (25, 265 and 26, 896, respectively), and their study included 650 of the MoBa individuals used in the current CFMR. However, the pre-pregnancy BMI instrument used by Tyrrell *et al.* was generated using the results from a GWAMA of 123, 865 individuals of European ancestry (Sardinians, Icelanders, and Estonians) (30). Furthermore, Tyrrell *et al.* analyzed 16 cohorts of European ancestry (from Europe, North America, and Australia). As recently pointed out by Zhang and colleagues (31), variations across populations may lead to biased estimates in an MR analysis. It is interesting to observe that our CFMR estimates were similar to the one-sample MR estimates (see Table 1 and Supplementary Figure 31). The fact that the CFMR estimates were similar to the observational association points to little confounding in the analyses. It is therefore expected that the one-sample MR estimates would be similar to the CFMR estimates (see Table 1 and Supplementary Figure 31). However, the standard errors of one-sample MR estimates are smaller than those of the CFMR estimates. These small standard errors are likely due to the inability of one-sample MR to handle overfitting, which results in an overly confident estimation (see Table 3.1 and Supplementary Figure 31) and biased inference when using one-sample MR. We illustrate this bias using simulations, and the results indicate that one-sample MR can be heavily biased even when the instrument is strong (*h*^2^ > 20%) (see Supplementary section 3.2 and Figure 32). CFMR, on the other hand, remains conservative even in the presence of weak instruments and strong confounding.

To conclude, we show that CFMR is a valuable new approach for MR analysis, particularly for small sample sizes and understudied exposures. It is especially useful for investigating exposures and outcomes that might be difficult or expensive to measure, or when dealing with populations made up of multiple ethnicities. Moreover, CFMR has the potential to enhance the power of two-sample MR in consortia-led meta-analyses, in which each cohort can apply CFMR to its study population and the results from each cohort subsequently meta-analyzed in the final step of the analysis. Our results showed that CFMR performed well when the sample is sufficiently large (≥ 100, 000) and even when the instruments are weak (*h*^2^ ≤ 1%). These advantageous features make CFMR an attractive tool to reassess the causal effect of poorly heritable traits, especially those with genotype data accessible through various public repositories, such as the Database of Genotypes and Phenotypes (dbGaP, https://www.ncbi.nlm.nih.gov/gap/) or the UK Biobank (https://www.ukbiobank.ac.uk/).

## Software

A typical CFMR run is provided as an R script at https://github.com/william-denault/CFMR. The scripts used for the current simulations and application to maternal pre-pregnancy BMI have also been deposited there.

## Acknowledgments

The authors thank Dr. Siri E. Håberg for providing access to the genotyped MoBa mother-child duos used in this study, and to Drs. Maria Magnus and Anders Skrondal for their insightful comments. Data were processed in digital labs at the HUNT Cloud, Norwegian University of Science and Technology, Trondheim, Norway. The authors are grateful to Max Denault for designing Figures 1 and The authors also thank all the MoBa participants, health professionals, laboratory technicians and field workers who have contributed to creating the MoBa dataset upon which this study is based. We thank the Norwegian Institute of Public Health (NIPH) for generating high-quality genomic data. This research is part of the HARVEST collaboration, supported by the Research Council of Norway (#229624). We also thank the NORMENT Centre for providing genotype data, funded by the Research Council of Norway (#223273), South East Norway Health Authority and KG Jebsen Stiftelsen. We further thank the Center for Diabetes Research, the University of Bergen for providing genotype data and performing quality control and imputation of the data funded by the ERC AdG project SELECTionPREDISPOSED, Stiftelsen Kristian Gerhard Jebsen, Trond Mohn Foundation, the Research Council of Norway, the Novo Nordisk Foundation, the University of Bergen, and the Western Norway health Authorities (Helse Vest).

## Supplementary material

## 1 2-fold CFMR

The first step in a 2-fold Cross-Fitting Mendelian randomization (CFMR) is the random partitioning of the original sample into two samples of equal size, *sample 1* and *sample 2*. After running a GWAS on each of these samples (*GWAS* _1_ and *GWAS* _2_) and clumping the results, SNPs that are suitable for use as IVs are selected and two sets of independent SNPs are created, *set*_1_ and *set*_2_. Using *sample 1* and *set*_1_, we build a predictor (*pred*_1_) of the exposure. Similarly, using *sample 2* and *set*_2_, we build a predictor (*pred*_2_) of the exposure. Next, we use *pred*_1_ to predict the exposure in *sample 2*. Using the predicted exposure as an instrument in *sample 2*, we estimate the causal effect of the exposure on the outcome in *sample 2* by performing a 2SLS. In a similar and complementary fashion, we estimate the causal effect of the exposure on the outcome in *sample 1* by performing a 2SLS using the predicted exposure in *sample 1* by *pred*_2_ as an instrument. The two estimates are referred to as 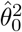 and 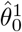, respectively. In this context, CFI refers to the use of the predicted exposure of *pred*_2_ as an instrument for *sample 1*, and vice versa (see Figure 1 in the main text).

Each instrument is then used to estimate the effect of the exposure on the outcome using a sample that does not overlap with the sample used to build the instrument, despite both samples stemming from the same source population. Thus, 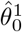 and 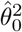 are unbiased two-sample MR estimates of the causal effect of the exposure on the outcome (1). Averaging these two estimates provides an unbiased estimate of the causal effect of the exposure on the outcome. The averaged estimate 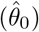 is referred to as the CFMR estimate of the exposure effect on the outcome. We refer to 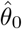 as the CFMR estimate of the exposure effect on the outcome.

## 2 Study description

The Norwegian Mother, father, and Child Cohort Study (MoBa) is an ongoing nationwide pregnancy cohort (2). Participants in MoBa were enrolled in the study between 1999 and 2008 from 50 of the 52 hospitals in Norway. The vast majority of MoBa participants are of Caucasian origin. The genotypes in the MoBa dataset were obtained from whole-blood DNA from parents and umbilical-cord blood DNA from newborns (3). We excluded stillbirths, twins, and children with missing data in the Medical Birth Registry of Norway (MBRN). Pre-pregnancy maternal BMI was calculated on the basis of self-reported height and weight. Information on the children’s birth weight was extracted from medical records.

Approximately 30, 000 mother-father-newborn trios in the MoBa dataset were genotyped using the Illumina HumanCoreExome BeadChip (San Diego, CA, USA), which contains more than 240,000 probes. We removed ethnic outliers based on visual checks using the first three principal components. We used the Haplotype Reference Consortium (HRC) reference data, version HRC.r1.1 (http://www.haplotype-reference-consortium.org/) for imputation of additional genotypes in the MoBa dataset. This was performed using the free genotype imputation and phasing service of the Sanger Imputation Server (https://imputation.sanger.ac.uk/).

The imputation quality was assessed by (i) hard-calling markers with an INFO quality score greater than 0.7, (ii) testing for Mendelian inconsistencies, excess of heterozygosity, and significant deviation from Hardy-Weinberg equilibrium (HWE), and (iii) screening for high rates of missingness. The remaining set of 7, 947, 894 SNPs met the following criteria and were included in the ten GWASes performed in the current CFMR: (i) call rate ≥ 98%, (ii) minor allele frequency (MAF) ≥ 1%, and (iii) HWE test P-value ≥ 10^4^. Samples with a call rate ≤ 98% and an excess heterozygosity ≥ 4*SD* were excluded. Finally, 1647 mother-child duos characterized by one or more of the following features were excluded: multiple births, stillbirths, congenital anomalies, births before 37 weeks gestation, pregnancy hypertension.

## 3 Simulations

### 3.1 CFMR unbiased estimation

Here, we show that CFMR behaves satisfactorily with respect to type I error, bias, and statistical power when LASSO is used to build the exposure predictors *pred*_1_ and *pred*_2_. We refer to CFMR as the estimator CFMR2. As explained in the manuscript, we chose CFMR2 over CFMR1 because, similarly to DML1 and DML2, CFMR2 exhibits a better finite sample size performance than CFMR1 (see (4)). Similar to the simulation set-up of Deng *et al.* (5), we consider a set of 300 independent variants (*V*_1_, …., *V*_300_) for each simulation, with each variant having a minor allele frequency of 0.3 and only the first five variants being associated with the exposure. The exposure is generated as 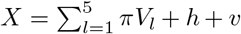 and the outcome is generated as *Y* = *β*_0_*X* + *h* + *u*, where *h* is a hidden confounder generated from a *N* (0, 2) distribution, and *v* and *u* are two correlated error terms generated from bivariate normal distribution.

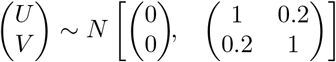

The variants have the same effect, denoted as *π*, where *π* is selected to ensure that the variants explain *h*^2^ = 20% of the variation in the exposure. Note that the only difference between our simulations and those of Deng *et al.* (5) is that the five variants associated with the exposure are assumed to be known in their simulations. Hence, Deng *et al.* only use the five truly-associated variants as instrument. In our case, we consider a more stringent set-up where we purposefully dilute the effects of the five truly-associated variants by adding 295 non-associated variants. This set-up makes it more challenging to construct accurate predictors of the exposure, particularly when the sample size is small and the IV is weak.

We applied CMFR to each simulated dataset using a LASSO-based IV and ten random splits. We considered various sample sizes (N = 1, 000 to 10, 000) and different *β*_0_ values (−0.08, −0.05, 0, 0.05, and 0.08), similar to the simulations by Deng *et al.* (5). For each combination of sample and effect size, we simulated 1000 datasets. Figure 1 summarizes the results of our simulations; we also provide a numerical summary of these simulations in Supplementary Tables 1 and 2. It is clear from Figure 1 that CFMR and two-sample MR have very similar power. CFMR also shows excellent control of the type I error for the different nominal levels tested (see Supplementary Table 1 and Supplementary Figure 4).

We also assessed the type I error, bias, and power of CFMR for different estimates of the variance explained by the exposure (*h*^2^ = 0%, 0.001%, 0.01%, 0.1%, 1%, 5%, and 10%) and different sample sizes (N = 1, 000, 5, 000, 10, 000, 50, 000, 100, 000, and 500, 000). For large sample sizes (N= 100, 000 or N=500, 000), we were unable to perform as many simulation as for smaller samples. Simulations were performed on a computer cluster with 32 CPUs and 128 GB RAM.

### 3.2 Comparison with one-sample MR

In this section, we perform a number of simulations to show that, under some settings where one-sample MR is heavily biased, CFMR remains conservative. The simulations were performed as follows. For each simulation, we consider a set of 300 independent variants (*V*_1_, …, *V*_300_), with each variant having a minor allele frequency of 0.3 and only the first five variants being associated with the exposure. The exposure is generated as 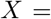 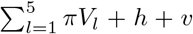 and the outcome is generated as *Y* = *β*_0_*X* + 40 × *h* + *u*, where *h* is a hidden confounder generated from a *N* (0, 2) distribution, and *v* and *u* are two correlated error terms generated from a bivariate normal distribution.

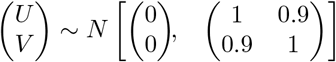

We consider the following two scenarios: 1) where *X* explains 10% of variance of *Y* (*h*^2^ = 10%), and 2) where *X* explains 20% of variance of *Y* (*h*^2^ = 20%). On each simulated dataset, we applied CFMR using a LASSO-based IV and ten random splits. We also build a predictor *pred_all_* of *X* using LASSO and the entire dataset. We then used the prediction *pred_all_* on the entire data as instrument. We refer to ‘one-sample MR estimates’ when we estimate the effect of *X* on *Y* using the prediction of *pred_all_*. We considered various sample sizes, ranging from 1, 000 to 50, 000, and *β*_0_ = 0.08. For each combination of sample and effect size, we simulated 1000 datasets. The results are summarized in Supplementary Table 7 and Figure 32.

## 4 Supplementary Material

### 4.1 Simulations results

**Figure 1:**
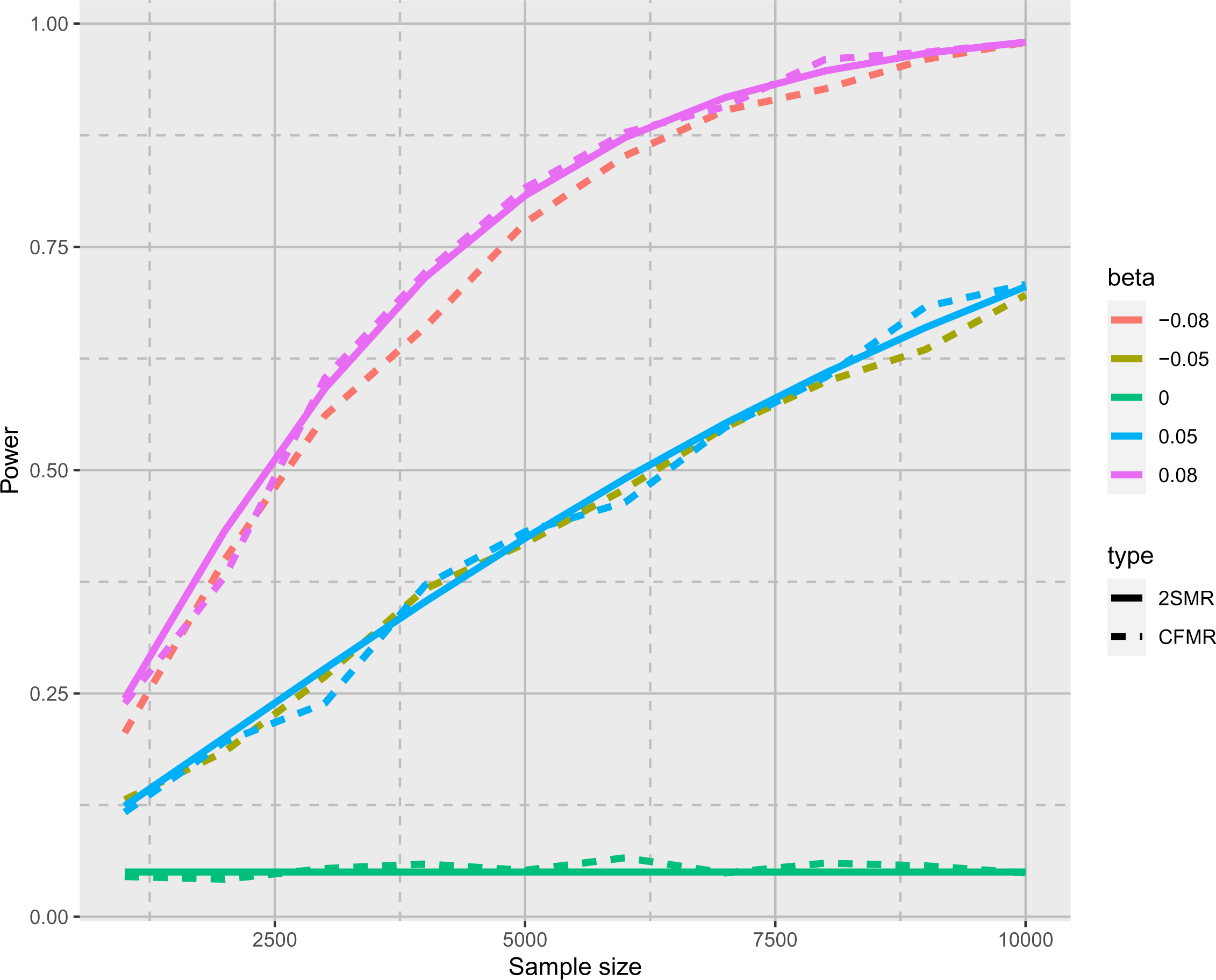
Power curves for CFMR and two-sample MR (2SMR) using the simulation set-up described in the Simulations section in the main text (with *h*^2^ = 20%). The dashed lines represent power curves for CFMR and the solid lines represent the theoretical power for 2SMR (5).

**Figure 2:**
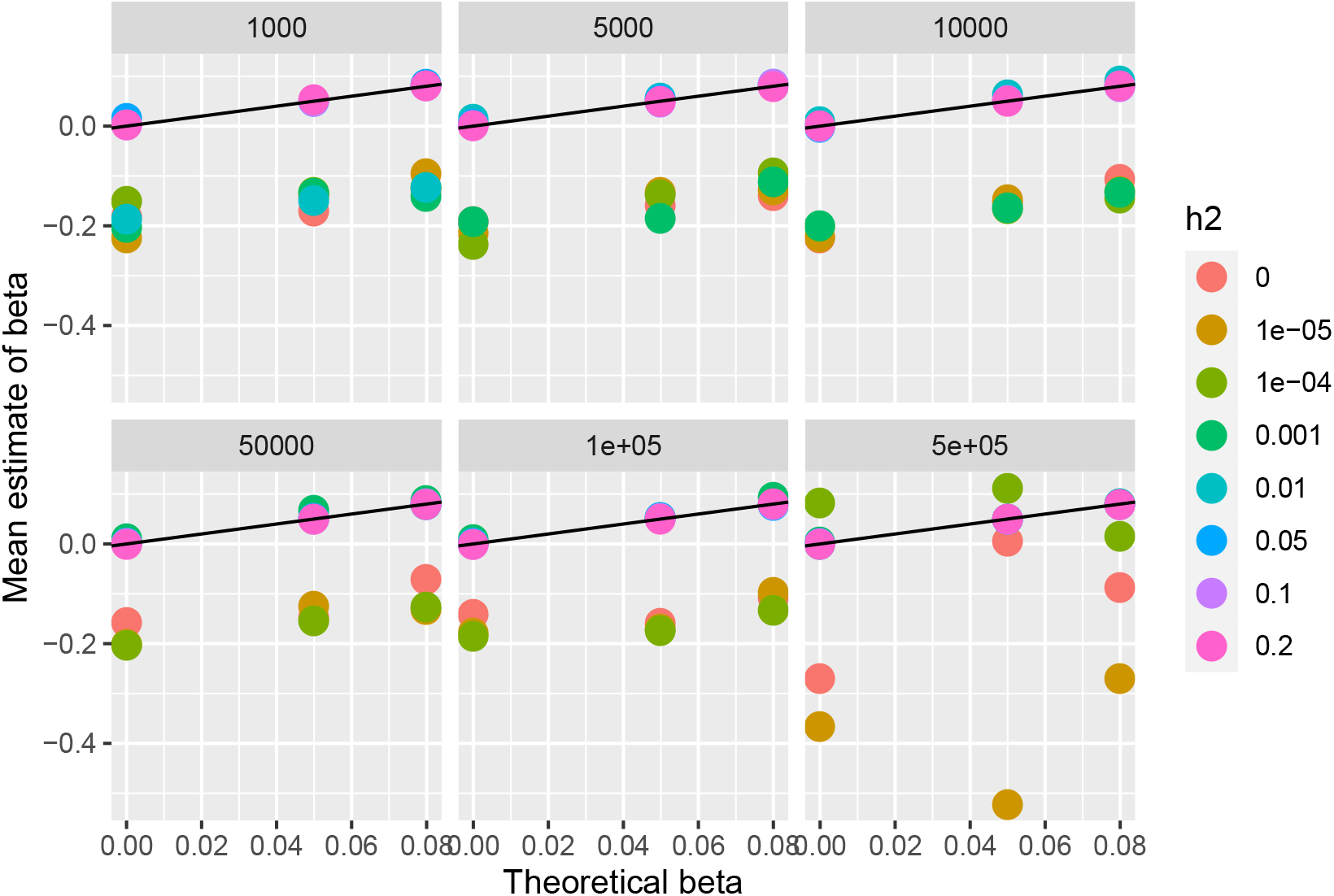
Mean estimate of beta by CFMR against the true beta for different values of beta, *h*^2^ and N.

**Figure 3:**
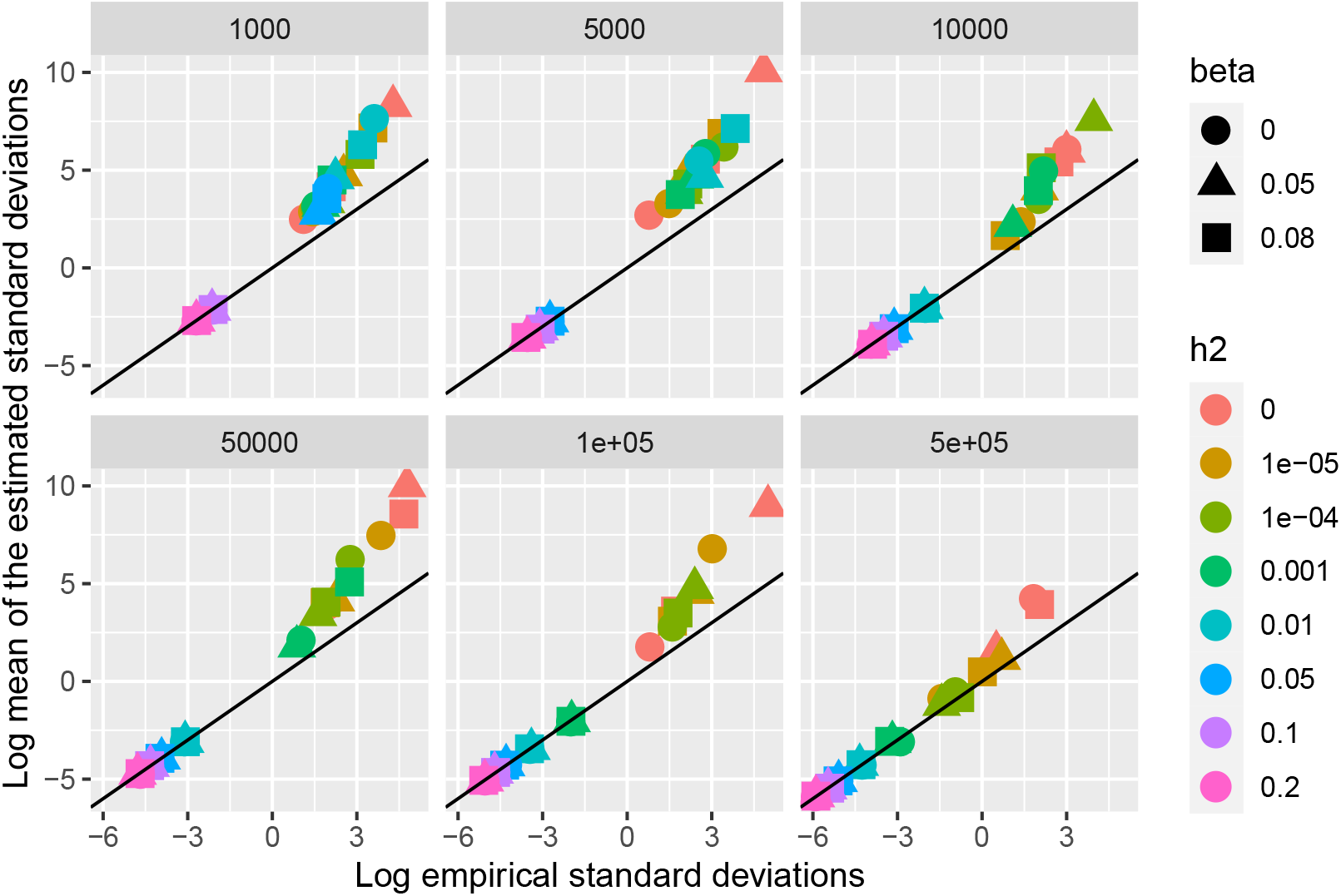
Empirical standard deviation of CFMR against the mean of the estimated standard deviations of CFMR for different values of beta, *h*^2^ and N.

**Figure 4:**
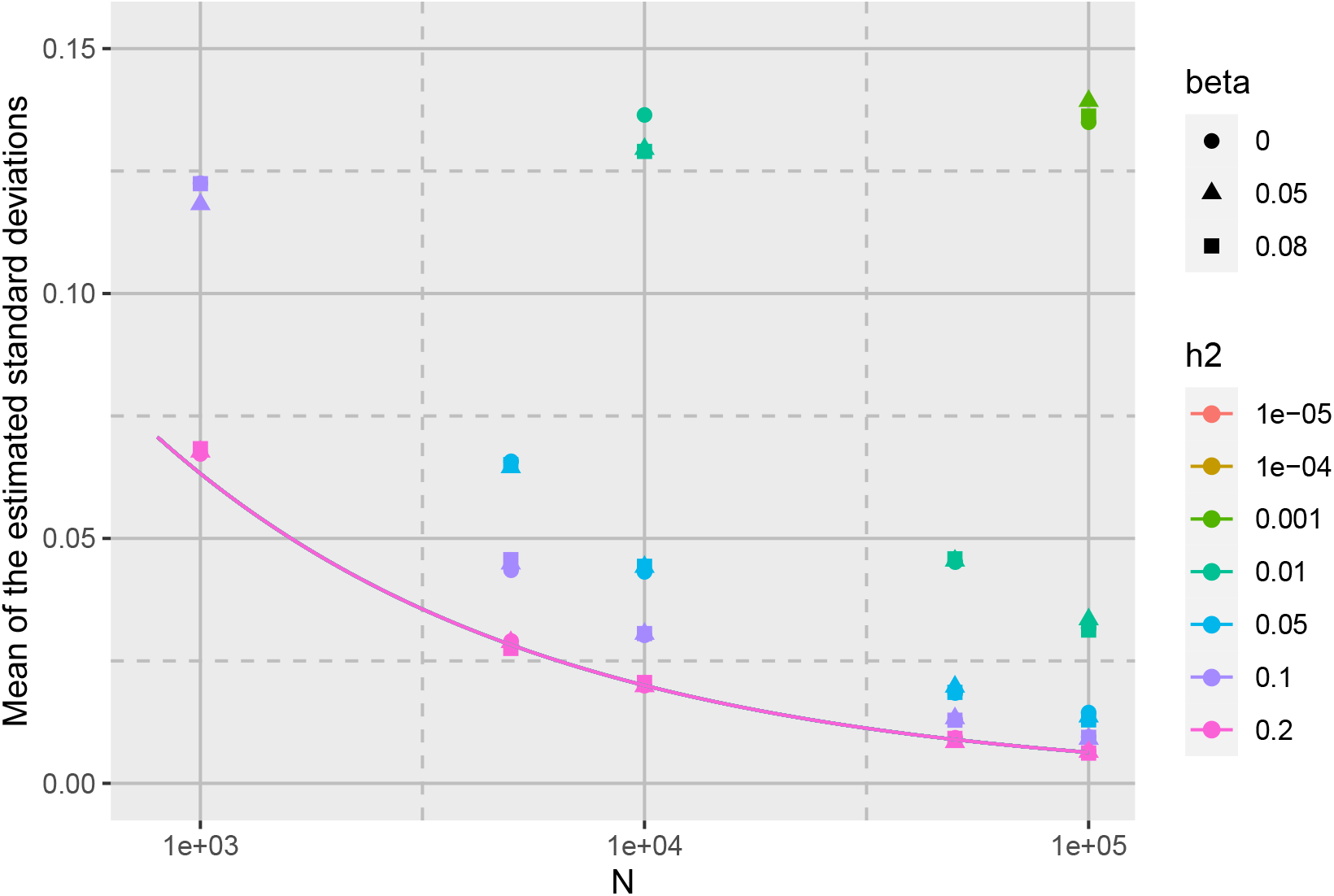
Mean of the estimated standard deviations of CFMR for different values of beta, *h*^2^ and N. Each configuration was simulated 1000 times. The solid line is the function 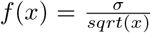, where *σ*^2^ is the variance of U in the simulations described in the Supplementary section 3.1.

**Figure 5:**
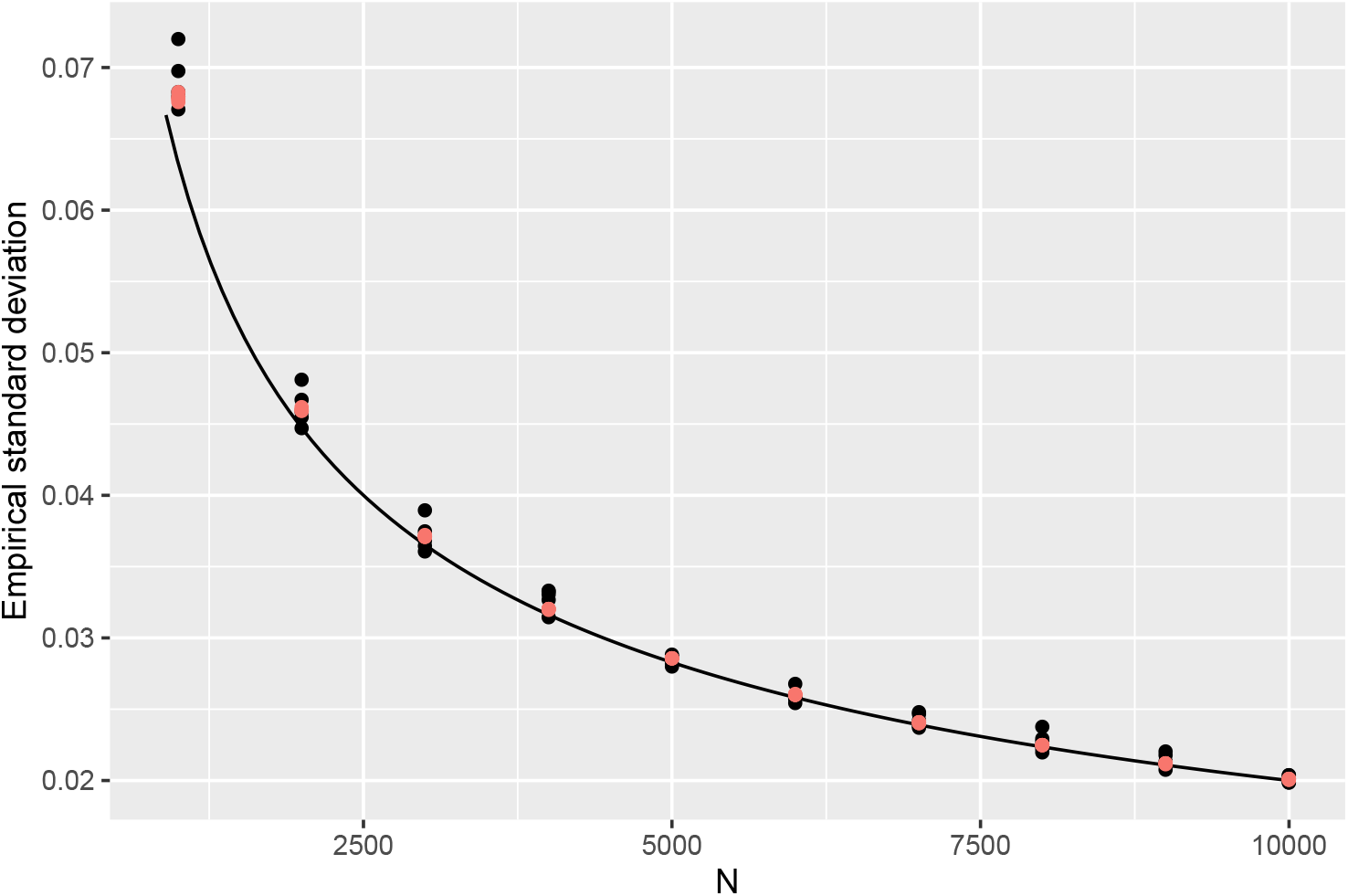
The black dots correspond to the empirical standard deviation of CFMR for various values of *θ*_0_. The red dots correspond to the empirical standard deviation of CFMR for various values of *θ*_0_ (*h*^2^ = 20%). The black line corresponds to the function 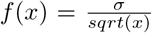, where *σ*^2^ is the variance of the simulations described in the Supplementary section 3.1.

**Figure 6:**
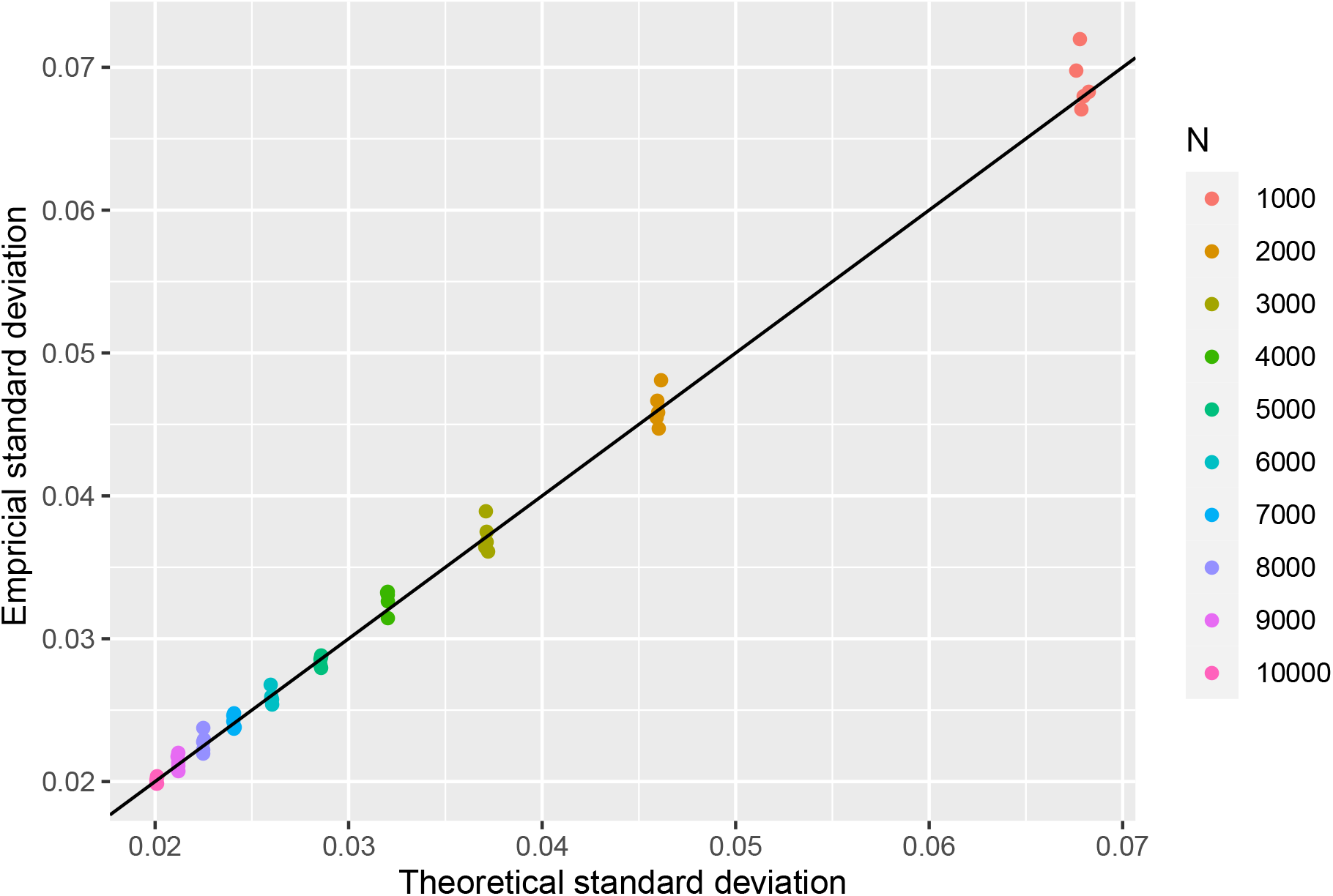
Empirical standard deviation of CFMR against the theoretical standard deviation of CFMR for different values of *θ*_0_ and N, with *h*^2^ = 20%.

**Figure 7:**
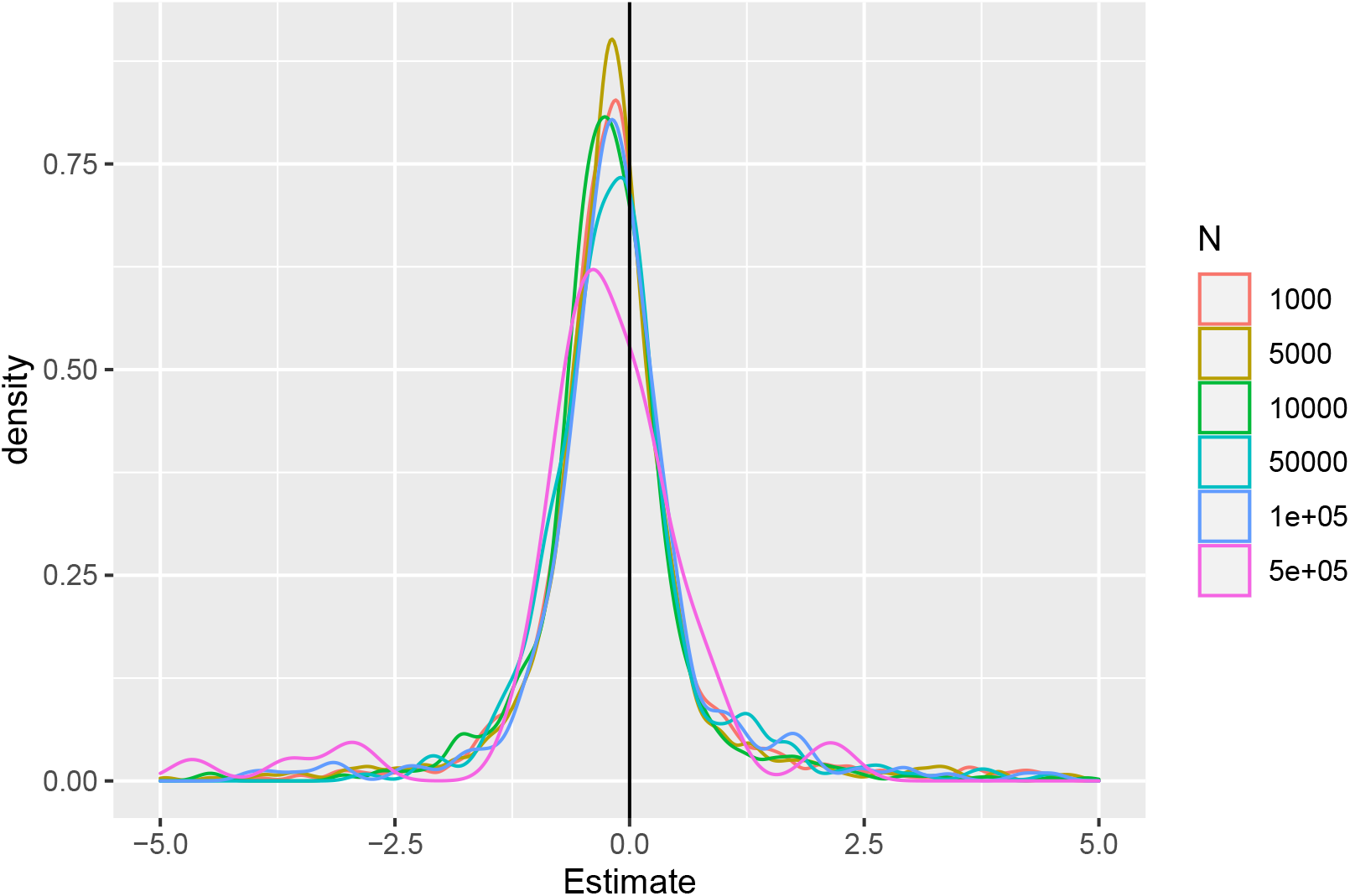
Density of the estimation of CFMR when no instrument is causally related to the exposure based on different sample sizes.

**Figure 8:**
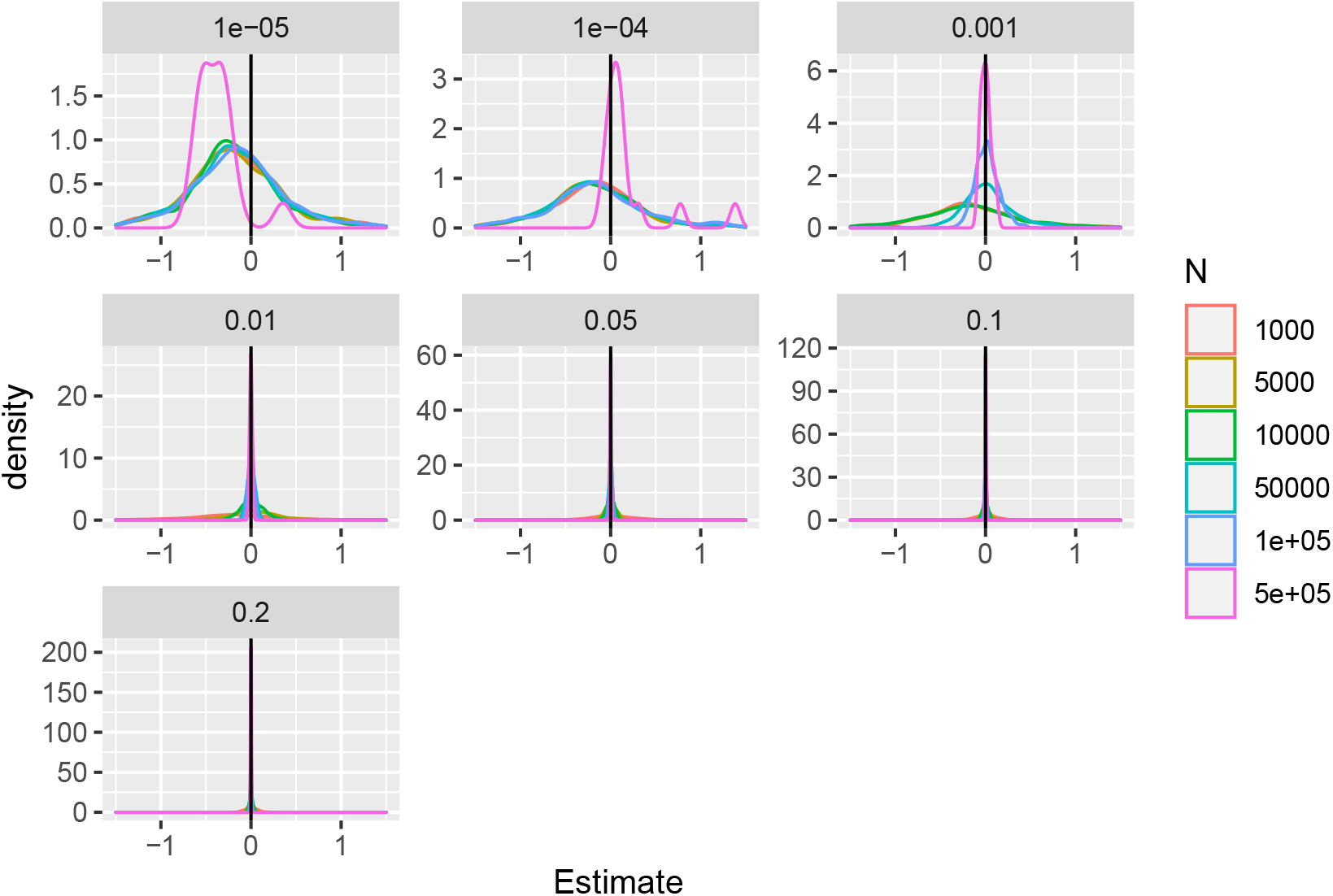
Density of the estimation of CFMR when *θ*_0_ = 0 for different sample sizes. The variance explained by the instrument (*h*^2^) is displayed on top of each plot.

**Table 1:** Estimated power of CFMR for different values of *β*_0_ and different sample sizes, with *h*^2^ = 20%.

| Sample size | $\beta_0$ | Mean($\hat{\beta}_0$) | sd($\hat{\beta}_0$) | Mean(sd($\hat{\beta}_0$)) | Power | Number of simulations <sup>1</sup> |
| --- | --- | --- | --- | --- | --- | --- |
| 1000 | -0.080 | -0.075 | 0.068 | 0.068 | 0.206 | 1000 |
| 1000 | -0.050 | -0.048 | 0.072 | 0.068 | 0.131 | 1000 |
| 1000 | 0.050 | 0.055 | 0.067 | 0.068 | 0.117 | 1000 |
| 1000 | 0.080 | 0.084 | 0.068 | 0.068 | 0.239 | 1000 |
| 2000 | -0.080 | -0.078 | 0.048 | 0.046 | 0.400 | 1000 |
| 2000 | -0.050 | -0.047 | 0.045 | 0.046 | 0.185 | 1000 |
| 2000 | 0.050 | 0.053 | 0.045 | 0.046 | 0.196 | 1000 |
| 2000 | 0.080 | 0.077 | 0.047 | 0.046 | 0.381 | 1000 |
| 3000 | -0.080 | -0.078 | 0.037 | 0.037 | 0.561 | 1000 |
| 3000 | -0.050 | -0.048 | 0.037 | 0.037 | 0.269 | 1000 |
| 3000 | 0.050 | 0.050 | 0.036 | 0.037 | 0.239 | 1000 |
| 3000 | 0.080 | 0.082 | 0.036 | 0.037 | 0.603 | 1000 |
| 4000 | -0.080 | -0.077 | 0.031 | 0.032 | 0.660 | 1000 |
| 4000 | -0.050 | -0.050 | 0.033 | 0.032 | 0.367 | 1000 |
| 4000 | 0.050 | 0.051 | 0.033 | 0.032 | 0.371 | 1000 |
| 4000 | 0.080 | 0.081 | 0.033 | 0.032 | 0.721 | 1000 |
| 5000 | -0.080 | -0.079 | 0.029 | 0.029 | 0.777 | 1000 |
| 5000 | -0.050 | -0.050 | 0.029 | 0.029 | 0.418 | 1000 |
| 5000 | 0.050 | 0.051 | 0.029 | 0.029 | 0.431 | 1000 |
| 5000 | 0.080 | 0.081 | 0.028 | 0.029 | 0.816 | 1000 |
| 6000 | -0.080 | -0.078 | 0.025 | 0.026 | 0.852 | 1000 |
| 6000 | -0.050 | -0.050 | 0.026 | 0.026 | 0.479 | 1000 |
| 6000 | 0.050 | 0.049 | 0.026 | 0.026 | 0.464 | 1000 |
| 6000 | 0.080 | 0.080 | 0.026 | 0.026 | 0.878 | 1000 |
| 7000 | -0.080 | -0.079 | 0.024 | 0.024 | 0.903 | 1000 |
| 7000 | -0.050 | -0.050 | 0.025 | 0.024 | 0.548 | 1000 |
| 7000 | 0.050 | 0.050 | 0.024 | 0.024 | 0.548 | 1000 |
| 7000 | 0.080 | 0.080 | 0.025 | 0.024 | 0.907 | 1000 |
| 8000 | -0.080 | -0.078 | 0.023 | 0.023 | 0.927 | 1000 |
| 8000 | -0.050 | -0.050 | 0.022 | 0.022 | 0.599 | 1000 |
| 8000 | 0.050 | 0.050 | 0.023 | 0.022 | 0.604 | 1000 |
| 8000 | 0.080 | 0.080 | 0.022 | 0.022 | 0.960 | 1000 |
| 9000 | -0.080 | -0.079 | 0.021 | 0.021 | 0.960 | 1000 |
| 9000 | -0.050 | -0.050 | 0.021 | 0.021 | 0.635 | 1000 |
| 9000 | 0.050 | 0.051 | 0.021 | 0.021 | 0.683 | 1000 |
| 9000 | 0.080 | 0.080 | 0.022 | 0.021 | 0.968 | 1000 |
| 10000 | -0.080 | -0.080 | 0.020 | 0.020 | 0.979 | 1000 |
| 10000 | -0.050 | -0.049 | 0.020 | 0.020 | 0.696 | 1000 |
| 10000 | 0.050 | 0.050 | 0.020 | 0.020 | 0.708 | 1000 |
| 10000 | 0.080 | 0.080 | 0.020 | 0.020 | 0.979 | 1000 |

<sup>1</sup>The column ‘Sample size’ corresponds to the sample size used in the simulation. ‘ $\beta_0$ ’ is the effect of $X$ on $Y$ to be estimated. ‘Mean $\hat{\beta}_0$ ’ corresponds to the average estimate of $\beta_0$ across simulations. ‘sd( $\hat{\beta}_0$ )’ corresponds to the observed standard deviation of $\hat{\beta}$ across simulations. ‘Mean (sd( $\hat{\beta}_0$ ))’ corresponds to the average of the estimated standard deviation of $\hat{\beta}_0$ . ‘Power’ corresponds to the proportion of the estimated P-value below 0.05. ‘Number of simulation’ is the number of simulations performed for the set of parameters (Sample size, $\beta_0$ , and $h^2$ in 3).

**Table 2:** Type I error of CFMR for different sample sizes, with *h*^2^ = 20%.

| Sample size | $\alpha$ level | | | Number of simulations |
| --- | --- | --- | --- | --- |
|  | 0.05 | 0.01 | 0.001 |  |
| 1000 | 0.045 | 0.011 | 0.001 | 1000 |
| 2000 | 0.042 | 0.008 | 0.001 | 1000 |
| 3000 | 0.054 | 0.014 | 0.003 | 1000 |
| 4000 | 0.059 | 0.013 | 0.001 | 1000 |
| 5000 | 0.052 | 0.007 | 0.000 | 1000 |
| 6000 | 0.066 | 0.015 | 0.002 | 1000 |
| 7000 | 0.049 | 0.006 | 0.001 | 1000 |
| 8000 | 0.060 | 0.010 | 0.001 | 1000 |
| 9000 | 0.057 | 0.014 | 0.002 | 1000 |
| 10000 | 0.049 | 0.010 | 0.001 | 1000 |

**Table 3:** Power of CFMR for different sample sizes and values of *h*^2^ (see the footnote of Table 1).

| $h^2$ | Sample size | $\beta$ | Mean $\hat{\beta}$ | sd($\hat{\beta}$) | Mean($\hat{sd}(\hat{\beta})$) | Power | Number of simulations |
| --- | --- | --- | --- | --- | --- | --- | --- |
| 0.00000 | 1000 | 0.05 | -0.1696 | 72.46 | 3997.26 | 0.019 | 1000 |
| 0.00000 | 1000 | 0.08 | -0.1268 | 8.09 | 63.37 | 0.010 | 1000 |
| 0.00000 | 5000 | 0.05 | -0.1581 | 126.03 | 23670.08 | 0.016 | 1000 |
| 0.00000 | 5000 | 0.08 | -0.1396 | 16.03 | 260.51 | 0.007 | 1000 |
| 0.00000 | 10000 | 0.05 | -0.1533 | 19.98 | 386.84 | 0.028 | 1000 |
| 0.00000 | 10000 | 0.08 | -0.1061 | 15.26 | 226.39 | 0.006 | 1000 |
| 0.00000 | 50000 | 0.05 | -0.1477 | 119.51 | 21979.87 | 0.012 | 335 |
| 0.00000 | 50000 | 0.08 | -0.0712 | 106.42 | 5236.63 | 0.006 | 330 |
| 0.00000 | 100000 | 0.05 | -0.1587 | 147.85 | 8001.65 | 0.012 | 335 |
| 0.00000 | 100000 | 0.08 | -0.1087 | 5.55 | 36.49 | 0.015 | 330 |
| 0.00000 | 500000 | 0.05 | 0.0058 | 1.65 | 4.67 | 0.030 | 67 |
| 0.00000 | 500000 | 0.08 | -0.0875 | 7.69 | 51.04 | 0.000 | 66 |
| 0.00001 | 100000 | 0.05 | -0.1981 | 2.87 | 27.43 | 0.007 | 150 |
| 0.00001 | 100000 | 0.08 | -0.2174 | 1.62 | 7.43 | 0.007 | 150 |
| 0.00010 | 100000 | 0.05 | -0.2185 | 35.57 | 1316.74 | 0.013 | 150 |
| 0.00010 | 100000 | 0.08 | -0.1148 | 2.22 | 5.30 | 0.013 | 150 |
| 0.00100 | 100000 | 0.05 | 0.0386 | 0.13 | 0.13 | 0.080 | 150 |
| 0.00100 | 100000 | 0.08 | 0.1057 | 0.13 | 0.13 | 0.140 | 150 |
| 0.01000 | 1000 | 0.05 | -0.1168 | 10.55 | 113.30 | 0.010 | 1000 |
| 0.01000 | 1000 | 0.08 | -0.1370 | 33.77 | 1006.66 | 0.014 | 1000 |
| 0.01000 | 5000 | 0.05 | 0.0520 | 21.79 | 196.09 | 0.020 | 1000 |
| 0.01000 | 5000 | 0.08 | 0.0852 | 65.17 | 2489.32 | 0.035 | 1000 |
| 0.01000 | 10000 | 0.05 | 0.0629 | 0.13 | 0.13 | 0.057 | 1000 |
| 0.01000 | 10000 | 0.08 | 0.0961 | 0.13 | 0.13 | 0.087 | 1000 |
| 0.01000 | 50000 | 0.05 | 0.0501 | 0.04 | 0.05 | 0.158 | 330 |
| 0.01000 | 50000 | 0.08 | 0.0730 | 0.05 | 0.05 | 0.385 | 335 |
| 0.01000 | 100000 | 0.05 | 0.0550 | 0.03 | 0.03 | 0.408 | 480 |
| 0.01000 | 100000 | 0.08 | 0.0802 | 0.03 | 0.03 | 0.707 | 485 |
| 0.01000 | 500000 | 0.05 | 0.0499 | 0.01 | 0.01 | 0.939 | 198 |
| 0.01000 | 500000 | 0.08 | 0.0817 | 0.01 | 0.01 | 1.000 | 201 |
| 0.05000 | 1000 | 0.05 | 0.0503 | 5.09 | 16.17 | 0.026 | 1000 |
| 0.05000 | 1000 | 0.08 | 0.0840 | 6.92 | 34.68 | 0.018 | 1000 |
| 0.05000 | 5000 | 0.05 | 0.0539 | 0.06 | 0.07 | 0.117 | 1000 |
| 0.05000 | 5000 | 0.08 | 0.0847 | 0.07 | 0.07 | 0.243 | 1000 |
| 0.05000 | 10000 | 0.05 | 0.0513 | 0.04 | 0.04 | 0.184 | 1000 |
| 0.05000 | 10000 | 0.08 | 0.0818 | 0.04 | 0.04 | 0.453 | 1000 |
| 0.05000 | 50000 | 0.05 | 0.0514 | 0.02 | 0.02 | 0.749 | 335 |

Table 3 – *Continued from previous page*
| $h^2$ | Sample size | $\beta$ | Mean $\hat{\beta}$ | sd($\hat{\beta}$) | Mean($\hat{sd}(\hat{\beta})$) | Power | Number of simulations |
| --- | --- | --- | --- | --- | --- | --- | --- |
| 0.05000 | 50000 | 0.08 | 0.0800 | 0.02 | 0.02 | 0.979 | 335 |
| 0.05000 | 100000 | 0.05 | 0.0529 | 0.01 | 0.01 | 0.958 | 335 |
| 0.05000 | 100000 | 0.08 | 0.0776 | 0.01 | 0.01 | 1.000 | 335 |
| 0.05000 | 500000 | 0.05 | 0.0495 | 0.01 | 0.01 | 1.000 | 67 |
| 0.05000 | 500000 | 0.08 | 0.0794 | 0.01 | 0.01 | 1.000 | 67 |
| 0.10000 | 1000 | 0.05 | 0.0497 | 0.12 | 0.12 | 0.047 | 1000 |
| 0.10000 | 1000 | 0.08 | 0.0817 | 0.12 | 0.12 | 0.095 | 1000 |
| 0.10000 | 5000 | 0.05 | 0.0486 | 0.04 | 0.04 | 0.210 | 1000 |
| 0.10000 | 5000 | 0.08 | 0.0838 | 0.05 | 0.04 | 0.485 | 1000 |
| 0.10000 | 10000 | 0.05 | 0.0515 | 0.03 | 0.03 | 0.374 | 1000 |
| 0.10000 | 10000 | 0.08 | 0.0799 | 0.03 | 0.03 | 0.760 | 1000 |
| 0.10000 | 50000 | 0.05 | 0.0508 | 0.01 | 0.01 | 0.958 | 335 |
| 0.10000 | 50000 | 0.08 | 0.0803 | 0.01 | 0.01 | 1.000 | 330 |
| 0.10000 | 100000 | 0.05 | 0.0509 | 0.01 | 0.01 | 1.000 | 335 |
| 0.10000 | 100000 | 0.08 | 0.0796 | 0.01 | 0.01 | 1.000 | 330 |
| 0.10000 | 500000 | 0.05 | 0.0500 | 0.00 | 0.00 | 1.000 | 201 |
| 0.10000 | 500000 | 0.08 | 0.0801 | 0.00 | 0.00 | 1.000 | 198 |
| 0.20000 | 1000 | 0.05 | 0.0527 | 0.07 | 0.07 | 0.114 | 1000 |
| 0.20000 | 1000 | 0.08 | 0.0816 | 0.07 | 0.07 | 0.233 | 1000 |
| 0.20000 | 5000 | 0.05 | 0.0505 | 0.03 | 0.03 | 0.420 | 1000 |
| 0.20000 | 5000 | 0.08 | 0.0799 | 0.03 | 0.03 | 0.809 | 1000 |
| 0.20000 | 10000 | 0.05 | 0.0499 | 0.02 | 0.02 | 0.705 | 1000 |
| 0.20000 | 10000 | 0.08 | 0.0805 | 0.02 | 0.02 | 0.981 | 1000 |
| 0.20000 | 50000 | 0.05 | 0.0495 | 0.01 | 0.01 | 1.000 | 330 |
| 0.20000 | 50000 | 0.08 | 0.0798 | 0.01 | 0.01 | 1.000 | 335 |
| 0.20000 | 100000 | 0.05 | 0.0505 | 0.01 | 0.01 | 1.000 | 330 |
| 0.20000 | 100000 | 0.08 | 0.0803 | 0.01 | 0.01 | 1.000 | 335 |
| 0.20000 | 500000 | 0.05 | 0.0497 | 0.00 | 0.00 | 1.000 | 132 |
| 0.20000 | 500000 | 0.08 | 0.0798 | 0.00 | 0.00 | 1.000 | 134 |

**Table 4:** Type I error of CFMR for different sample sizes and values of *h*^2^.

| $h^2$ | Sample size | $\alpha$ level | | | Number of simulations |
| --- | --- | --- | --- | --- | --- |
|  |  | 0.05 | 0.01 | 0.001 |  |
| 0.00000 | 1000 | 0.0380 | 0.0000 | 0.0000 | 1000 |
| 0.00000 | 5000 | 0.0240 | 0.0010 | 0.0000 | 1000 |
| 0.00000 | 10000 | 0.0270 | 0.0050 | 0.0010 | 1000 |
| 0.00000 | 50000 | 0.0239 | 0.0090 | 0.0000 | 335 |
| 0.00000 | 100000 | 0.0209 | 0.0000 | 0.0000 | 335 |
| 0.00000 | 500000 | 0.0299 | 0.0000 | 0.0000 | 67 |
| 0.00001 | 1000 | 0.0260 | 0.0020 | 0.0000 | 1000 |
| 0.00001 | 5000 | 0.0210 | 0.0030 | 0.0000 | 1000 |
| 0.00001 | 10000 | 0.0280 | 0.0020 | 0.0000 | 1000 |
| 0.00001 | 50000 | 0.0250 | 0.0040 | 0.0000 | 1000 |
| 0.00001 | 100000 | 0.0333 | 0.0067 | 0.0000 | 150 |
| 0.00010 | 1000 | 0.0270 | 0.0040 | 0.0000 | 1000 |
| 0.00010 | 5000 | 0.0250 | 0.0010 | 0.0000 | 1000 |
| 0.00010 | 10000 | 0.0290 | 0.0040 | 0.0000 | 1000 |
| 0.00010 | 50000 | 0.0200 | 0.0020 | 0.0000 | 1000 |
| 0.00010 | 100000 | 0.0333 | 0.0000 | 0.0000 | 150 |
| 0.00100 | 1000 | 0.0270 | 0.0010 | 0.0000 | 1000 |
| 0.00100 | 5000 | 0.0250 | 0.0020 | 0.0000 | 1000 |
| 0.00100 | 10000 | 0.0300 | 0.0040 | 0.0000 | 1000 |
| 0.00100 | 50000 | 0.0230 | 0.0010 | 0.0000 | 1000 |
| 0.00100 | 100000 | 0.0800 | 0.0200 | 0.0000 | 150 |
| 0.01000 | 1000 | 0.0240 | 0.0020 | 0.0000 | 2000 |
| 0.01000 | 5000 | 0.0230 | 0.0030 | 0.0005 | 2000 |
| 0.01000 | 10000 | 0.0445 | 0.0075 | 0.0000 | 2000 |
| 0.01000 | 50000 | 0.0487 | 0.0067 | 0.0007 | 1335 |
| 0.01000 | 100000 | 0.0722 | 0.0206 | 0.0021 | 485 |
| 0.01000 | 500000 | 0.0597 | 0.0100 | 0.0000 | 201 |
| 0.05000 | 1000 | 0.0260 | 0.0040 | 0.0000 | 1000 |
| 0.05000 | 5000 | 0.0430 | 0.0110 | 0.0020 | 1000 |
| 0.05000 | 10000 | 0.0480 | 0.0080 | 0.0010 | 1000 |
| 0.05000 | 50000 | 0.0273 | 0.0061 | 0.0000 | 330 |
| 0.05000 | 100000 | 0.0697 | 0.0182 | 0.0000 | 330 |
| 0.05000 | 500000 | 0.0758 | 0.0000 | 0.0000 | 66 |
| 0.10000 | 1000 | 0.0560 | 0.0050 | 0.0000 | 1000 |
| 0.10000 | 5000 | 0.0410 | 0.0080 | 0.0010 | 1000 |
| 0.10000 | 10000 | 0.0470 | 0.0140 | 0.0010 | 1000 |
| 0.10000 | 50000 | 0.0388 | 0.0060 | 0.0000 | 335 |
| 0.10000 | 100000 | 0.0537 | 0.0119 | 0.0000 | 335 |
| 0.10000 | 500000 | 0.0398 | 0.0000 | 0.0000 | 201 |
| 0.20000 | 1000 | 0.0400 | 0.0100 | 0.0010 | 1000 |
| 0.20000 | 5000 | 0.0510 | 0.0120 | 0.0010 | 1000 |
| 0.20000 | 10000 | 0.0490 | 0.0120 | 0.0030 | 1000 |
| 0.20000 | 50000 | 0.0657 | 0.0119 | 0.0000 | 335 |
| 0.20000 | 100000 | 0.0657 | 0.0090 | 0.0000 | 335 |
| 0.20000 | 500000 | 0.0522 | 0.0075 | 0.0000 | 134 |

## 5 Estimating the effect of pre-pregnancy maternal BMI on fetal birth weight using CFMR

**Figure 9:**
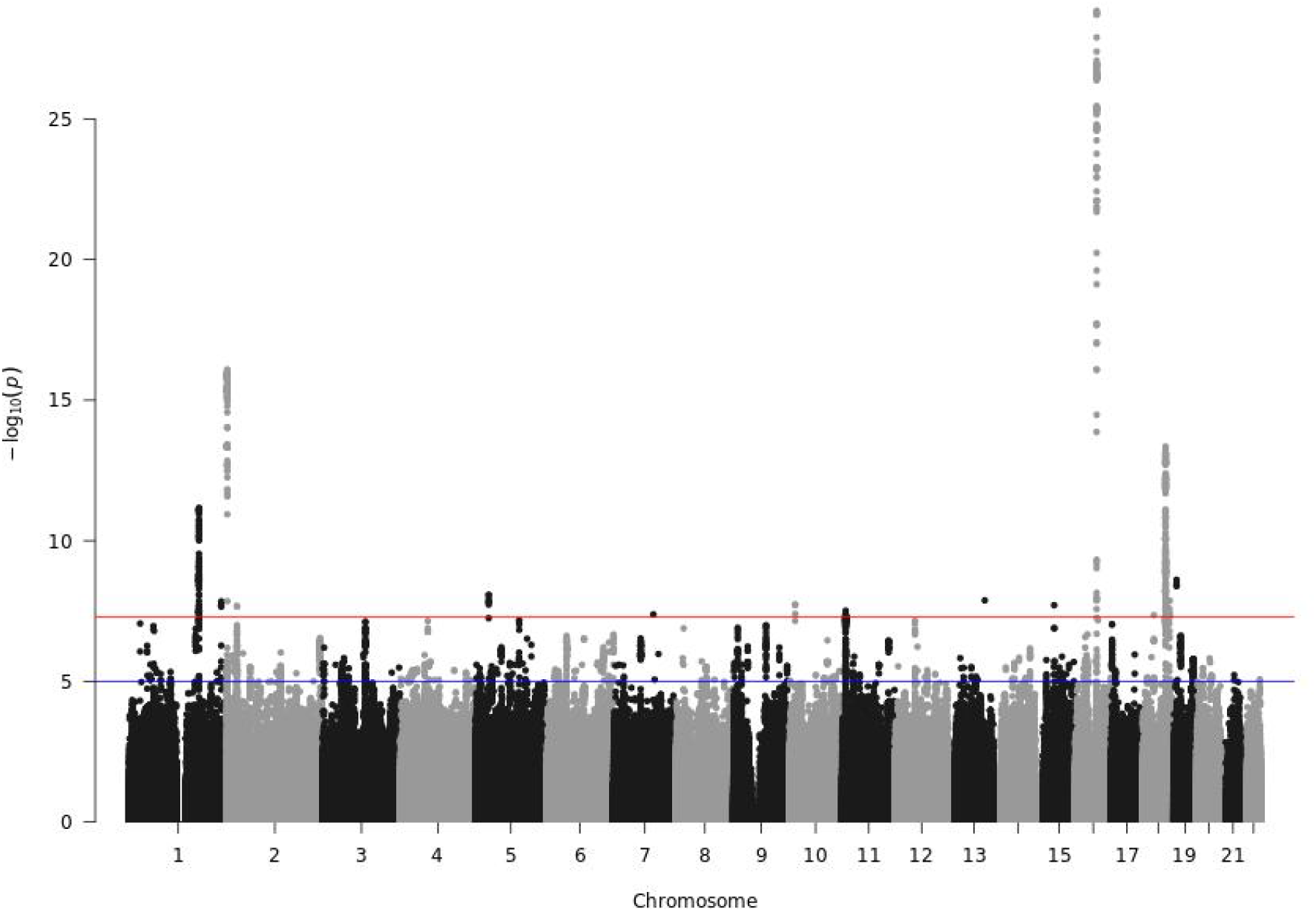
Manhattan plot from the GWAS performed on the first split number 1. The horizontal red line corresponds to the genome-wide significance threshold 5 × 10^−8^. The horizontal blue line corresponds to a significance threshold of 10^−5^.

**Figure 10:**
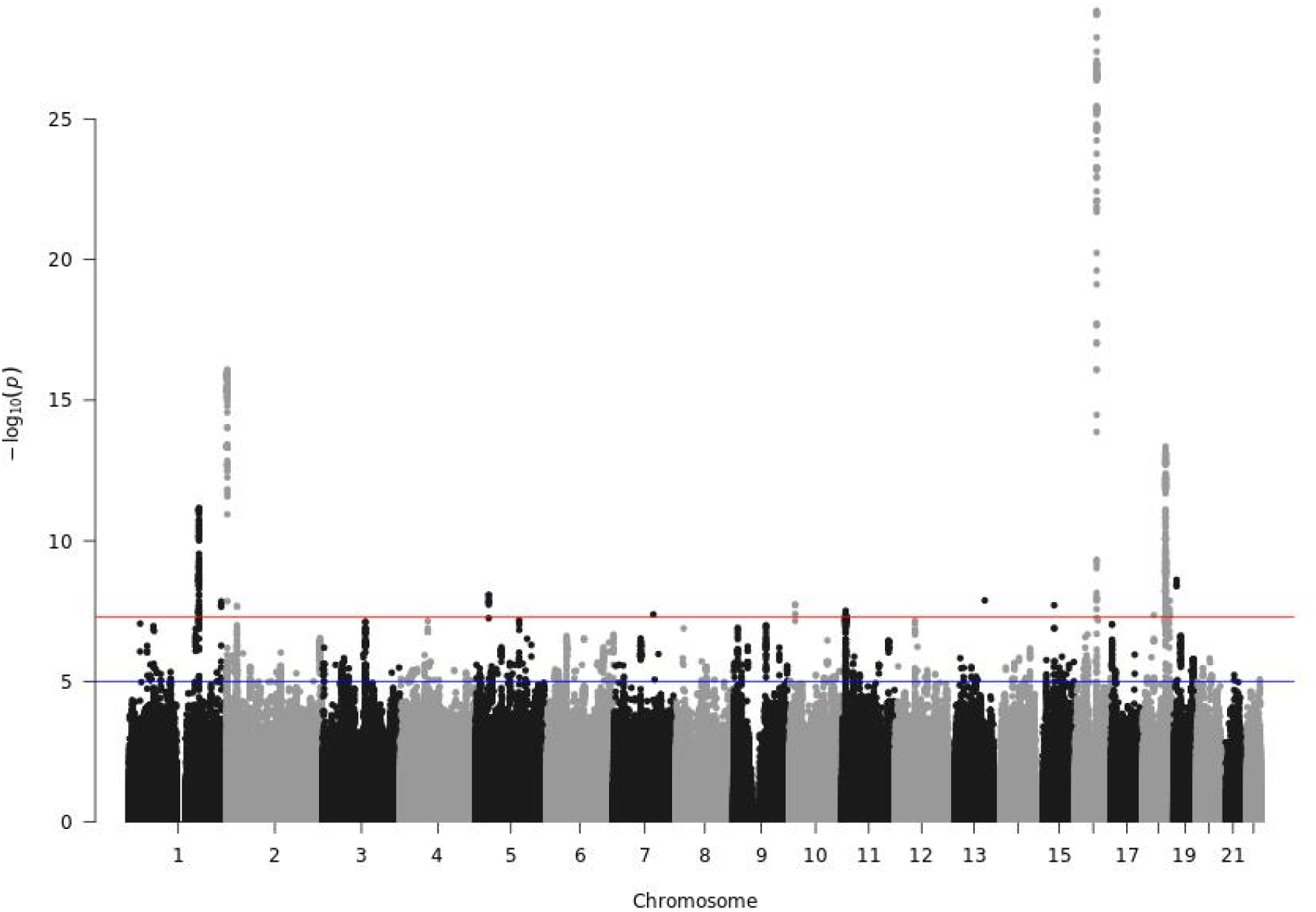
Manhattan plot from the GWAS performed on split number 2. The horizontal red line corresponds to the genome-wide significance threshold 5 × 10^−8^. The horizontal blue line corresponds to a significance threshold of 10^−5^.

**Figure 11:**
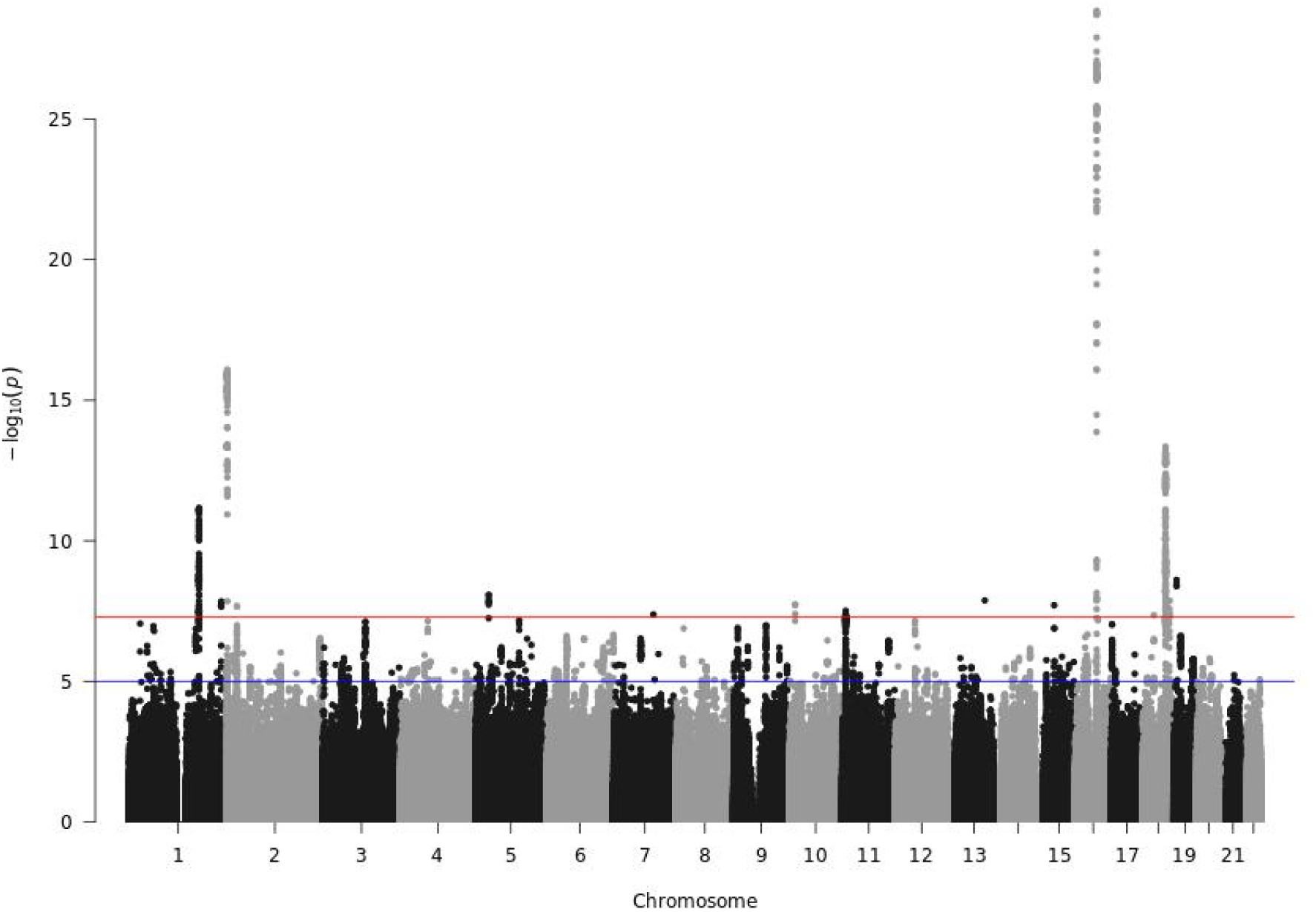
Manhattan plot from the GWAS performed on split number 3. The horizontal red line corresponds to the genome-wide significance threshold 5 × 10^−8^. The horizontal blue line corresponds to a significance threshold of 10^−5^.

**Figure 12:**
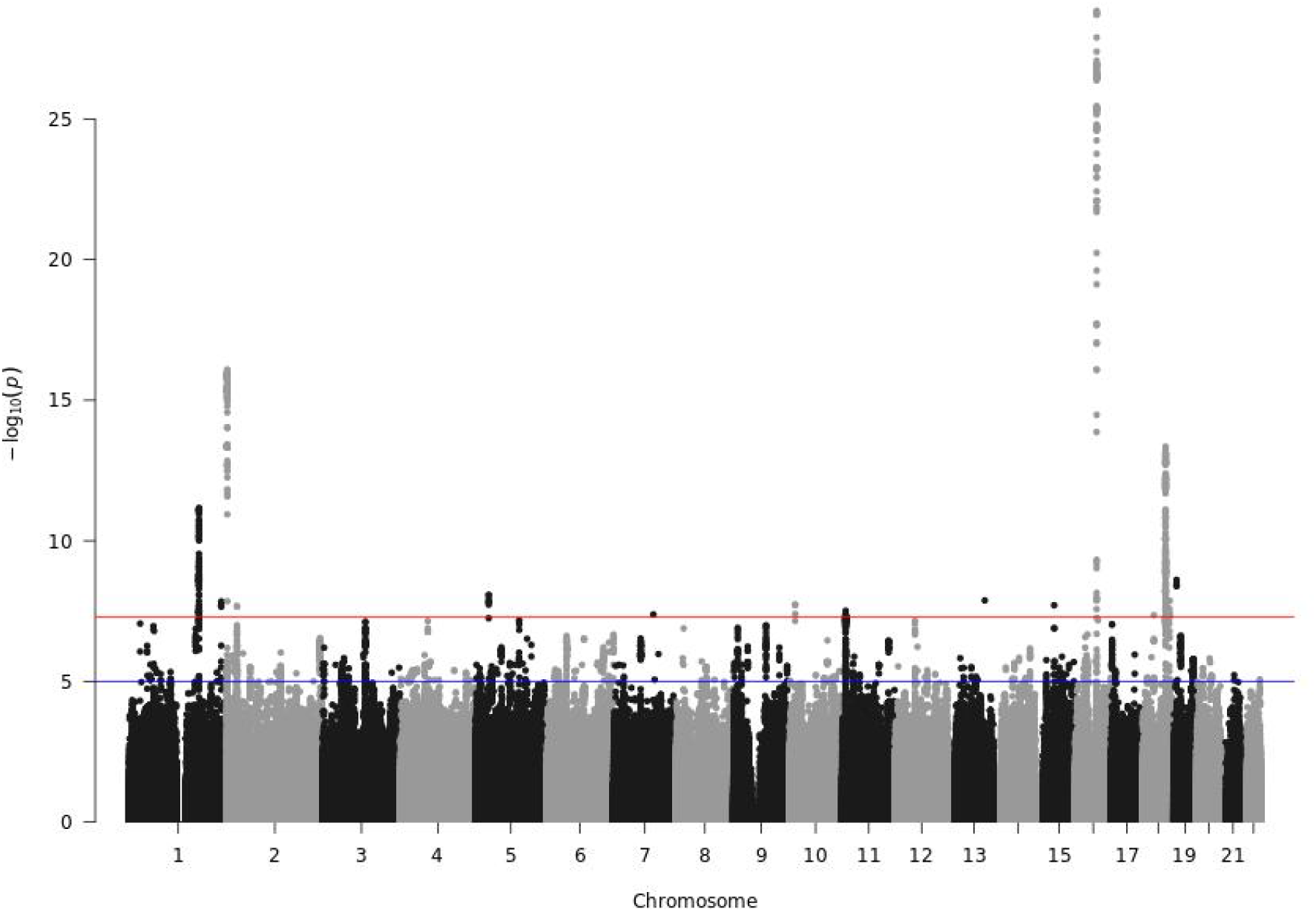
Manhattan plot from the GWAS performed on split number 4. The horizontal red line corresponds to the genome-wide significance threshold 5 × 10^−8^. The horizontal blue line corresponds to a significance threshold of 10^−5^.

**Figure 13:**
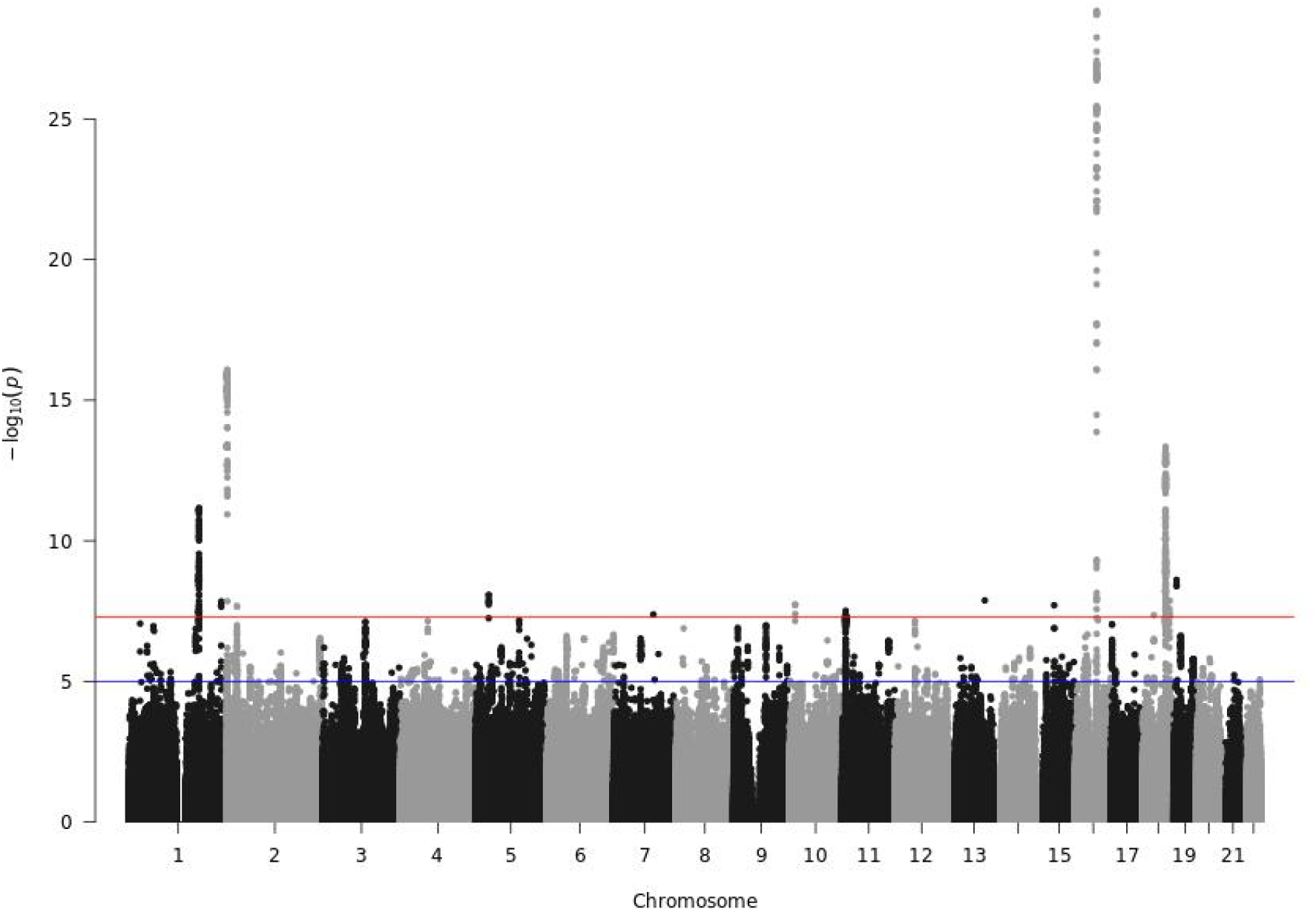
Manhattan plot from the GWAS performed on split number 5. The horizontal red line corresponds to the genome-wide significance threshold 5 × 10^−8^. The horizontal blue line corresponds to a significance threshold of 10^−5^.

**Figure 14:**
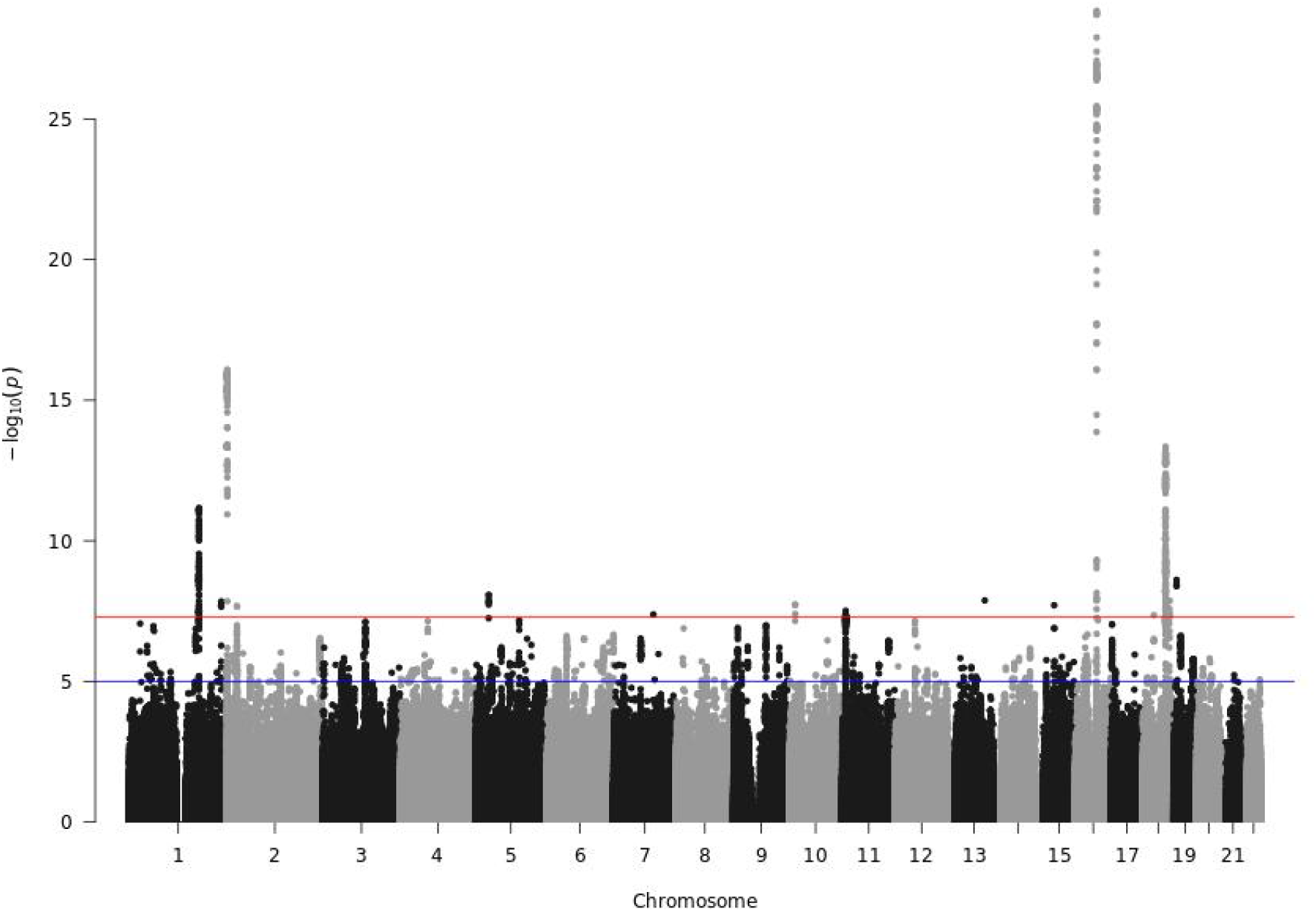
Manhattan plot from the GWAS performed on split number 6. The horizontal red line corresponds to the genome-wide significance threshold 5 × 10^−8^. The horizontal blue line corresponds to a significance threshold of 10^−5^.

**Figure 15:**
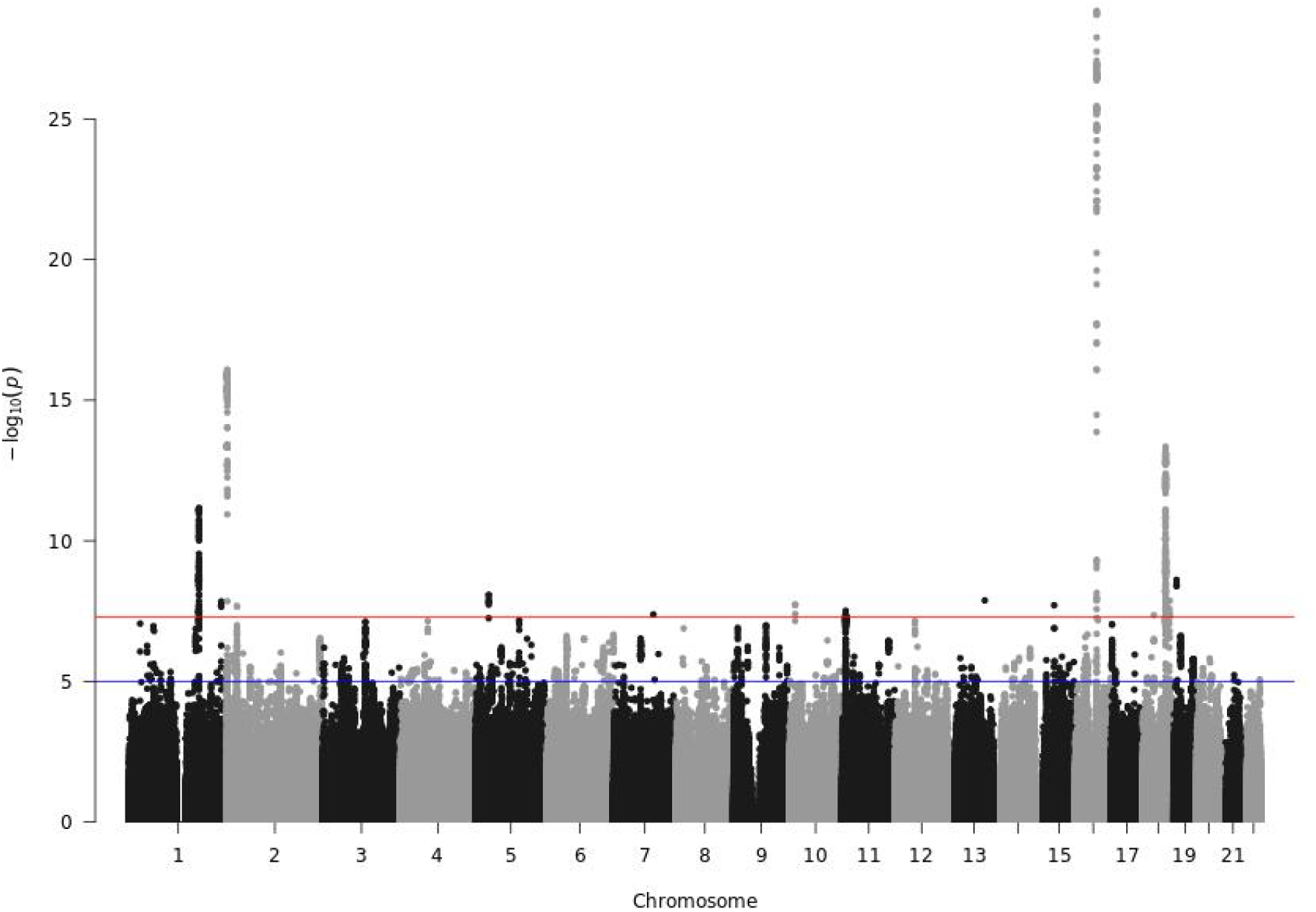
Manhattan plot from the GWAS performed on split number 7. The horizontal red line corresponds to the genome-wide significance threshold 5 × 10^−8^. The horizontal blue line corresponds to a significance threshold of 10^−5^.

**Figure 16:**
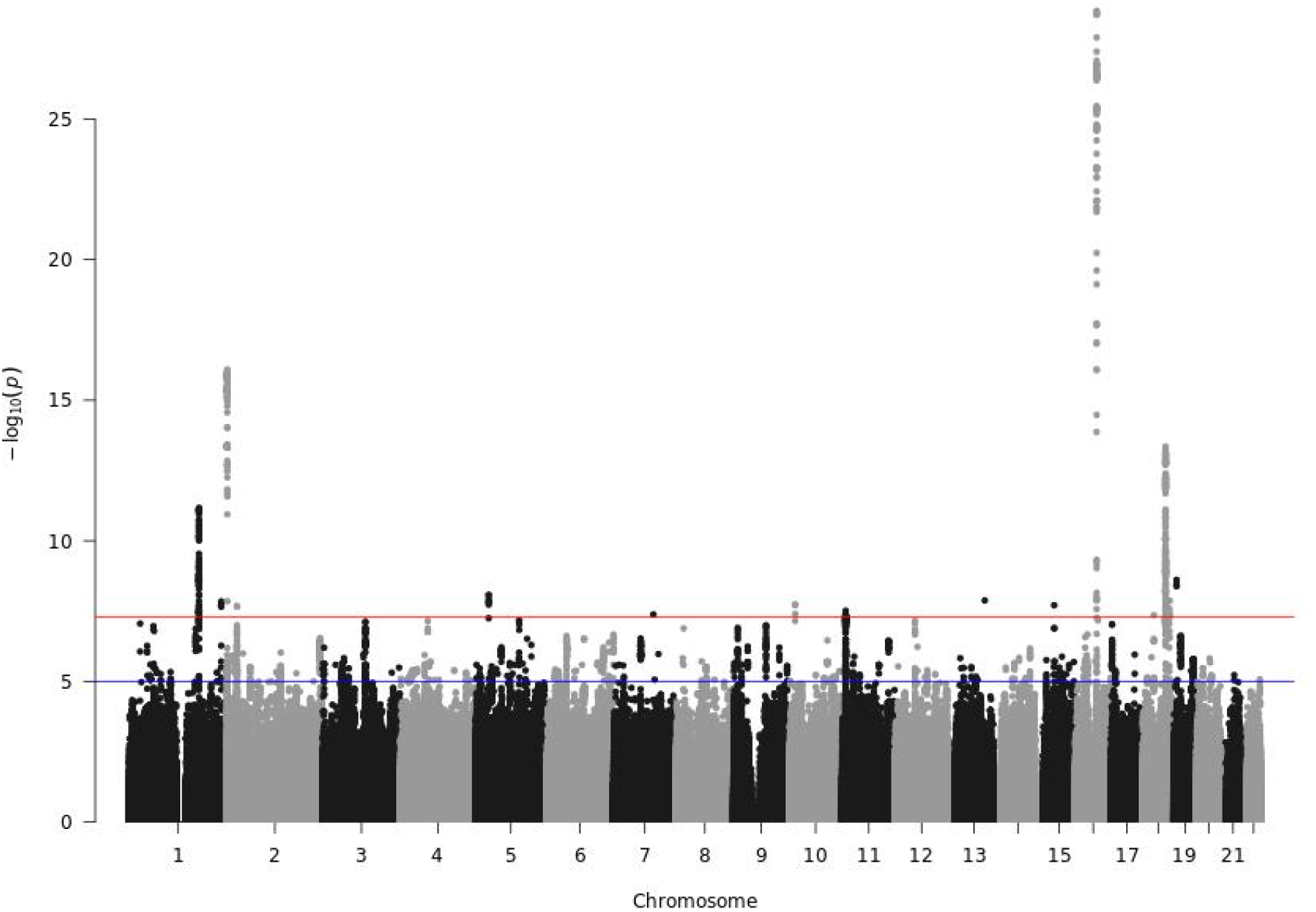
Manhattan plot from the GWAS performed on split number 8. The horizontal red line corresponds to the genome-wide significance threshold 5 × 10^−8^. The horizontal blue line corresponds to a significance threshold of 10^−5^.

**Figure 17:**
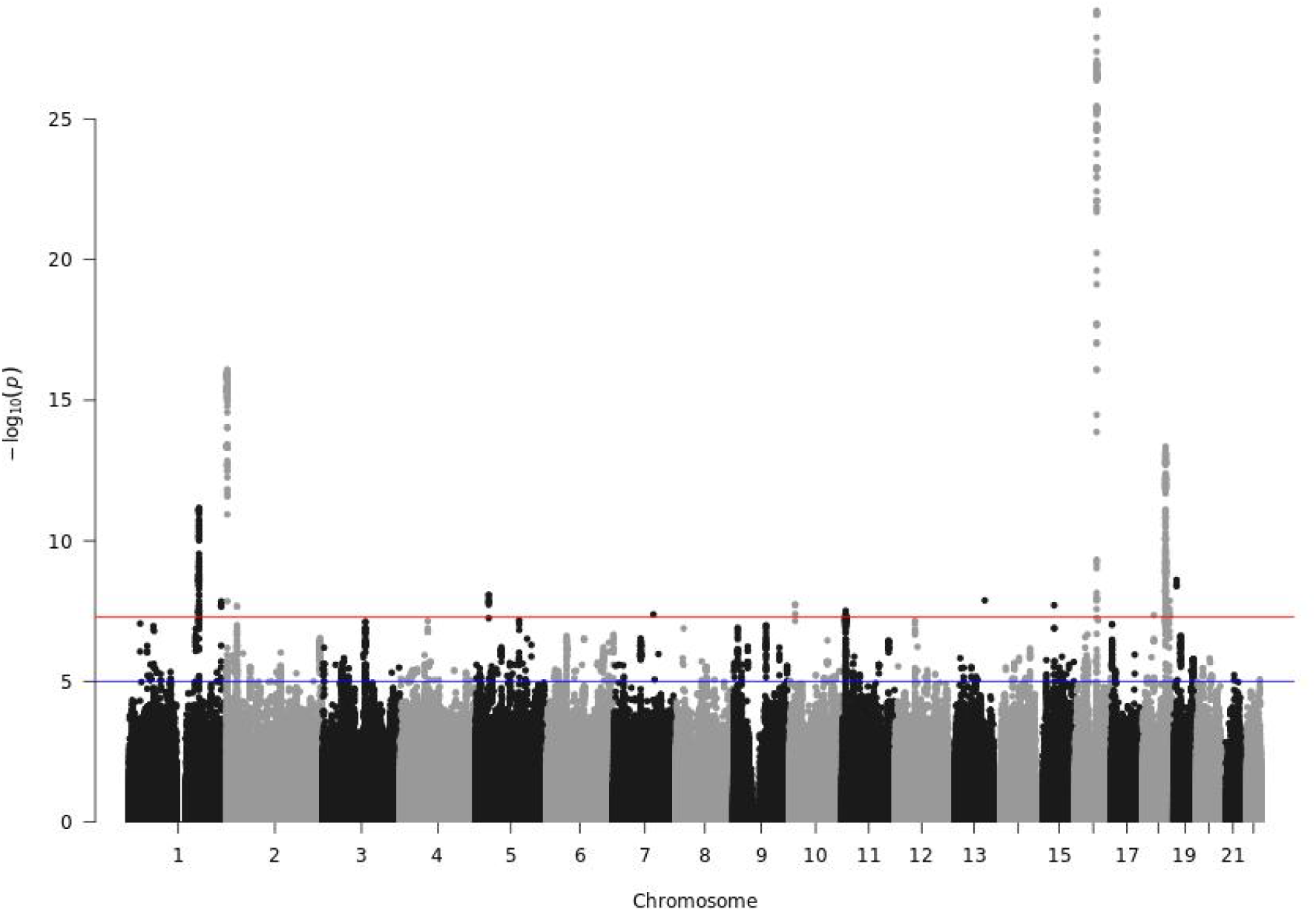
Manhattan plot from the GWAS performed on split number 9. The horizontal red line corresponds to the genome-wide significance threshold 5 × 10^−8^. The horizontal blue line corresponds to a significance threshold of 10^−5^.

**Figure 18:**
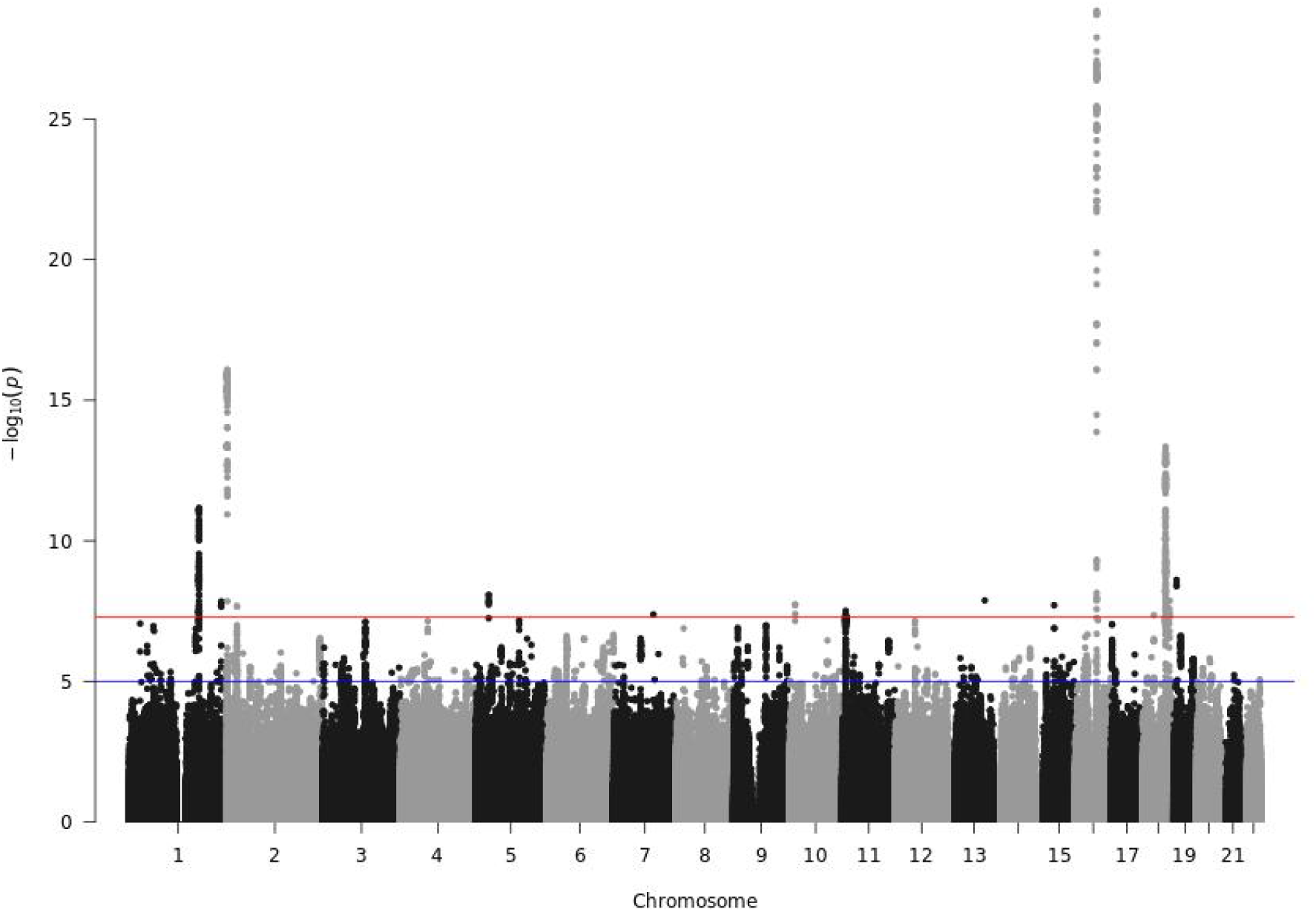
Manhattan plot from the GWAS performed on split number 10. The horizontal red line corresponds to the genome-wide significance threshold 5 × 10^−8^. The horizontal blue line corresponds to a significance threshold of 10^−5^.

**Figure 19:**
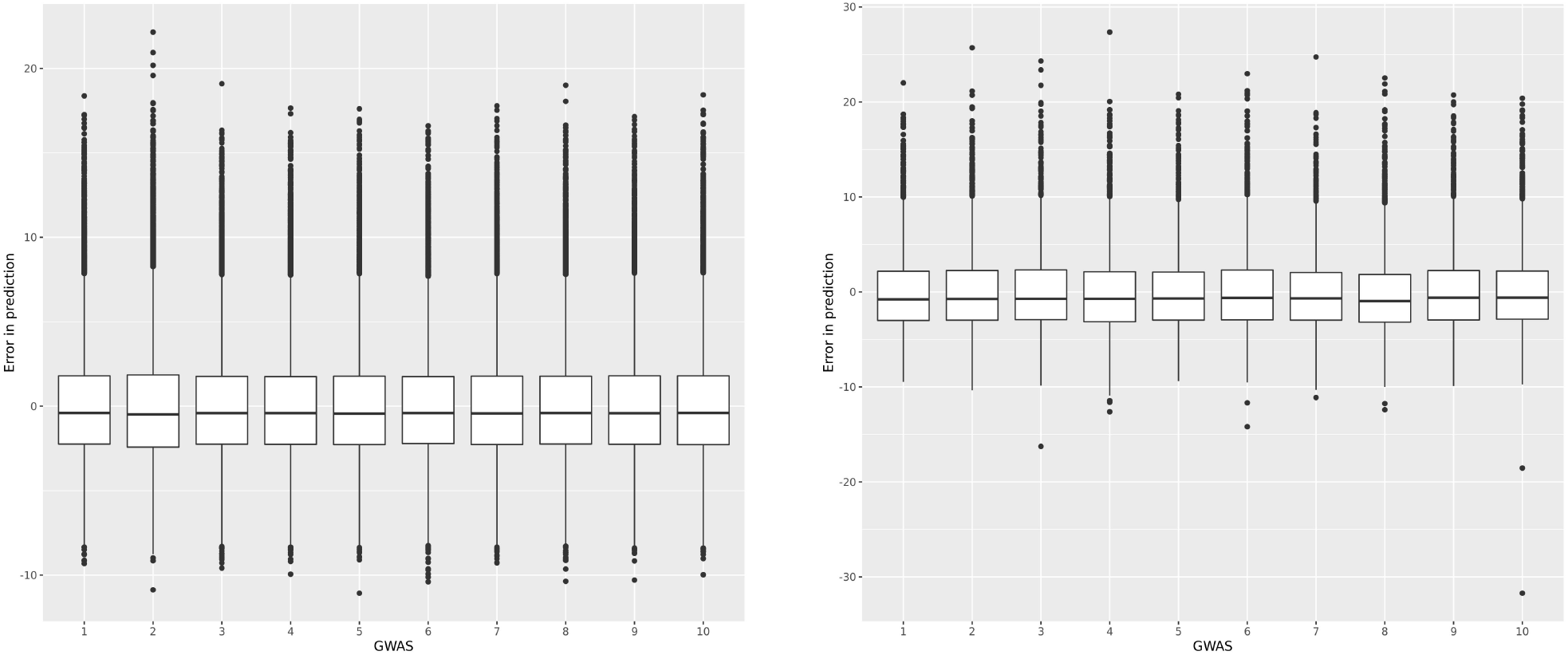
Predicted pre-pregnancy BMI performance on test and training sets. Left panel, the boxplot of the difference between the predicted pre-pregnancy BMI and the observed pre-pregnancy BMI on each training set, using a P-value threshold of 10^−3^. Right panel, the boxplot of the difference between the predicted pre-pregnancy BMI and the observed pre-pregnancy BMI on each test set, using a P-value threshold of 10^−3^.

**Figure 20:**
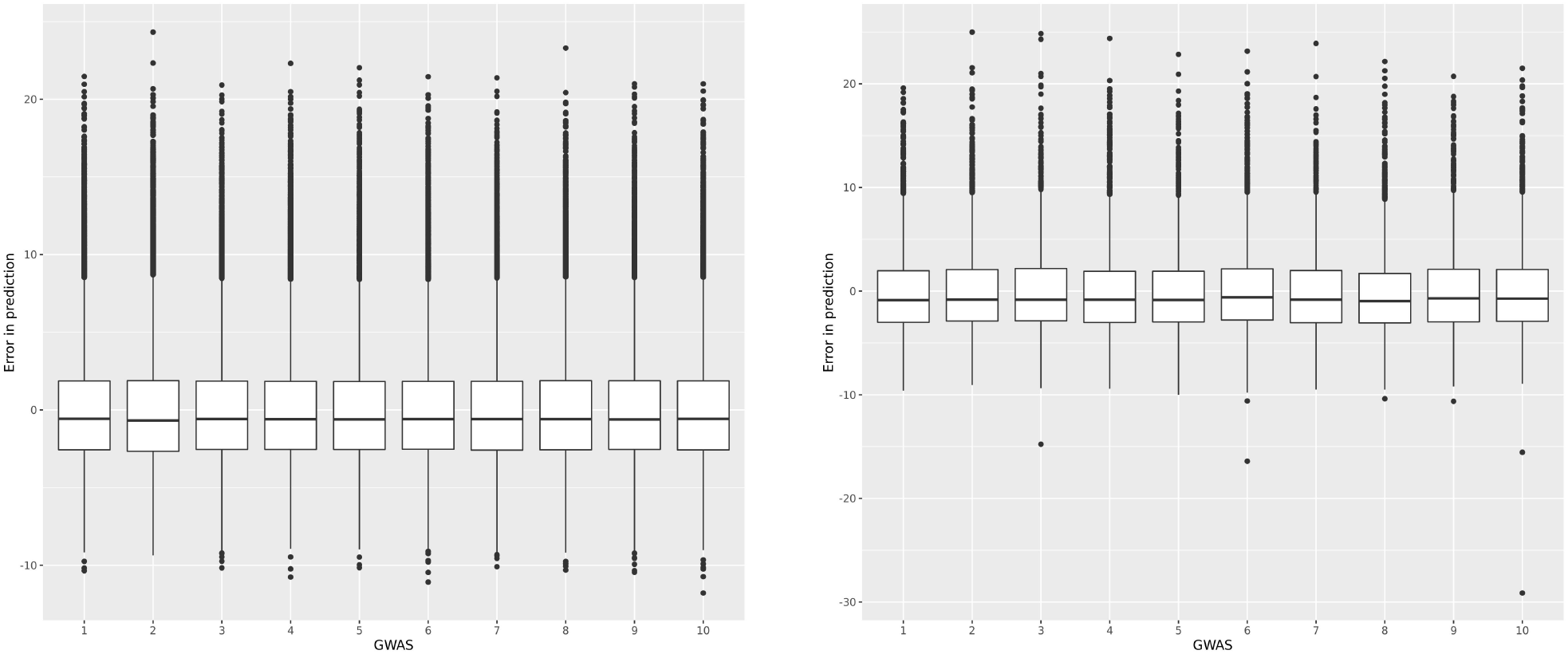
Predicted pre-pregnancy BMI performance on test and training sets. Left panel, the boxplot of the difference between the predicted pre-pregnancy BMI and the observed pre-pregnancy BMI on each training set, using a P-value threshold of 10^−4^. Right panel, the boxplot of the difference between the predicted pre-pregnancy BMI and the observed pre-pregnancy BMI on each test set, using a P-value threshold of 10^−4^.

**Figure 21:**
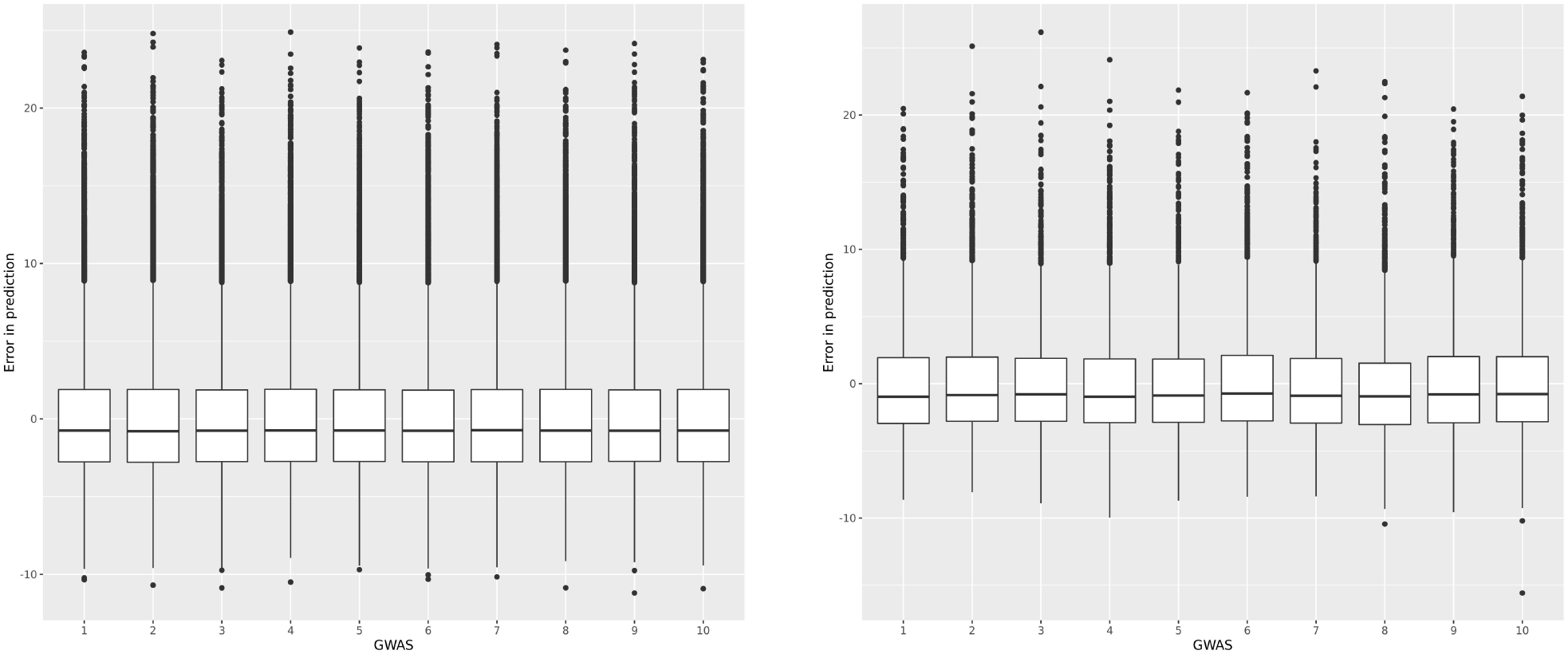
Predicted pre-pregnancy BMI performance on test and training sets. Left panel, the boxplot of the difference between the predicted pre-pregnancy BMI and the observed pre-pregnancy BMI on each training set, using a P-value threshold of 10^−5^. Right panel, the boxplot of the difference between the predicted pre-pregnancy BMI and the observed pre-pregnancy BMI on each test set, using a P-value threshold of 10^−5^.

**Figure 22:**
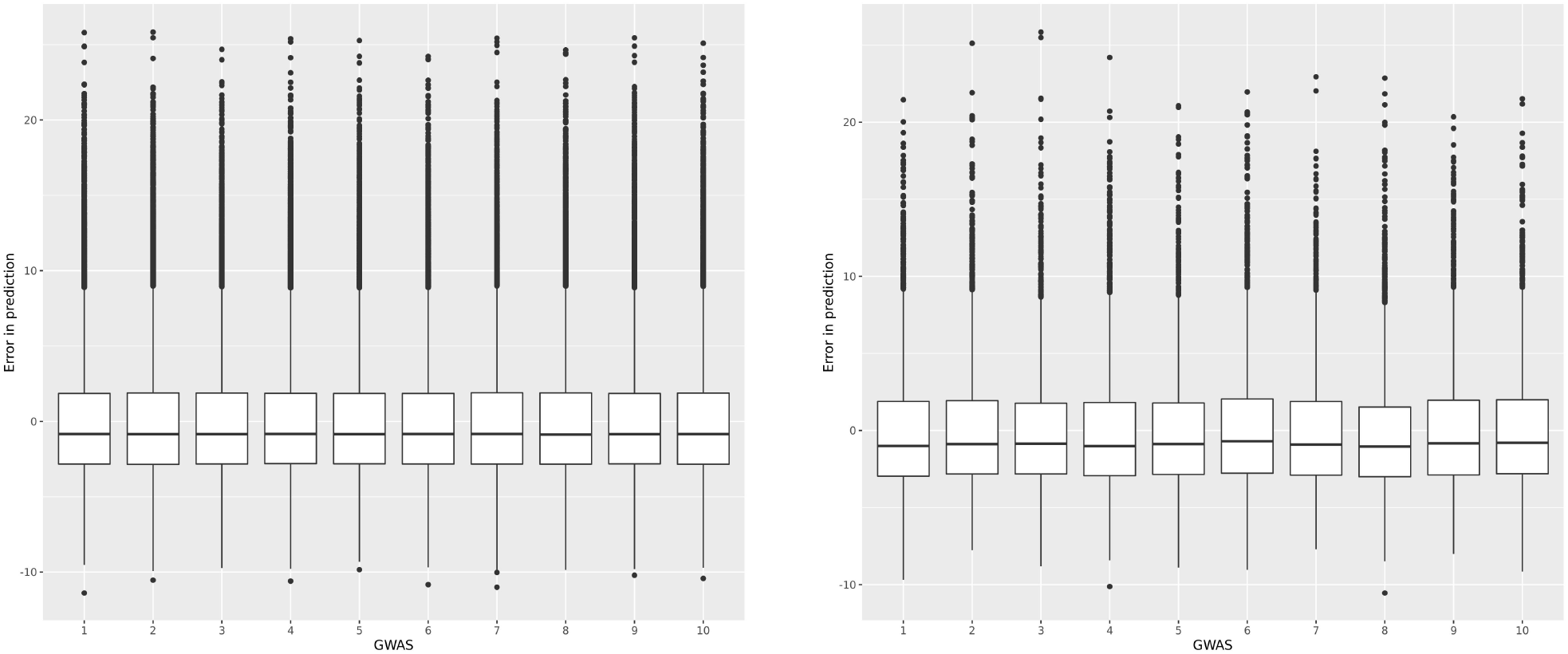
Predicted pre-pregnancy BMI performance on test and training sets. Left panel, the boxplot of the difference between the predicted pre-pregnancy BMI and the observed pre-pregnancy BMI on each training set, using a P-value threshold of 10^−6^. Right panel, the boxplot of the difference between the predicted pre-pregnancy BMI and the observed pre-pregnancy BMI on each test set, using a P-value threshold of 10^−6^.

**Figure 23:**
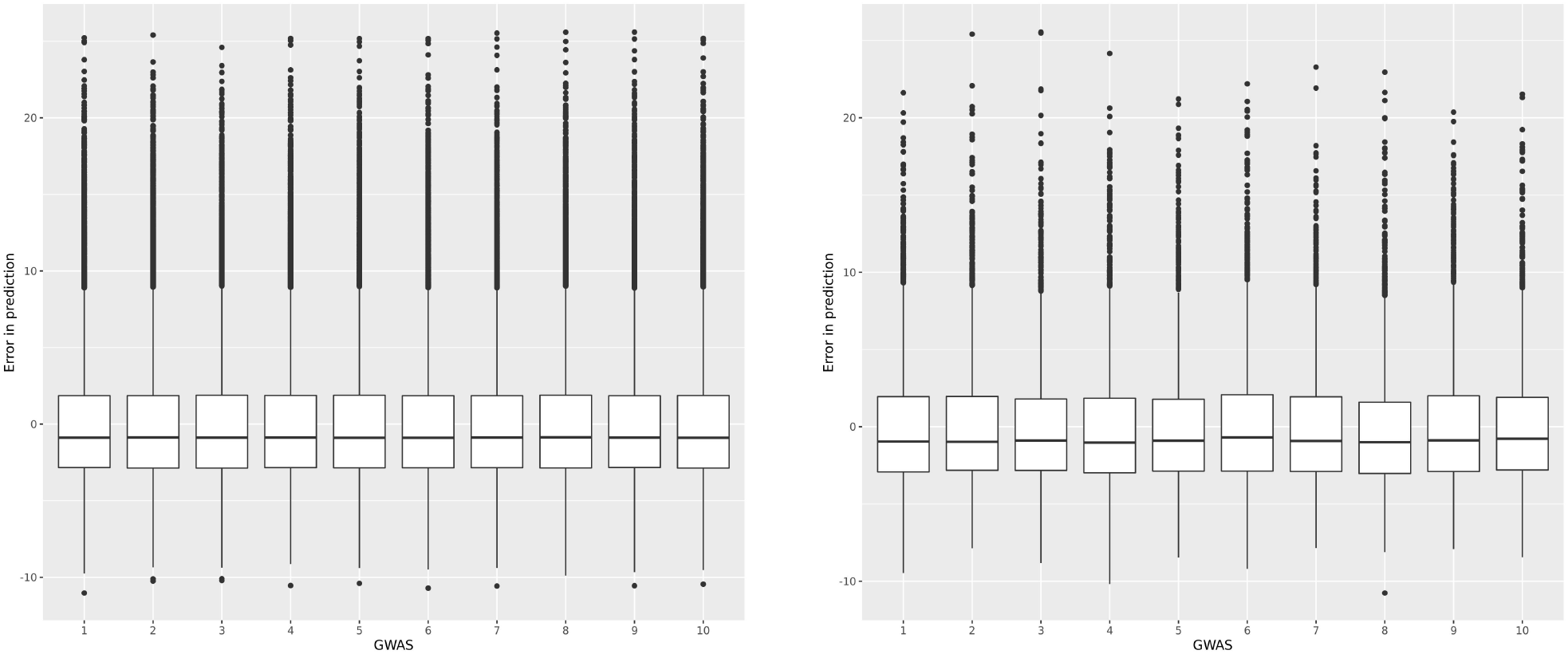
Predicted pre-pregnancy BMI performance on test and training sets. Left panel, the boxplot of the difference between the predicted pre-pregnancy BMI and the observed pre-pregnancy BMI on each training set, using a P-value threshold of 10^−7^. Right panel, the boxplot of the difference between the predicted pre-pregnancy BMI and the observed pre-pregnancy BMI on each test set, using a P-value threshold of 10^−7^.

**Figure 24:**
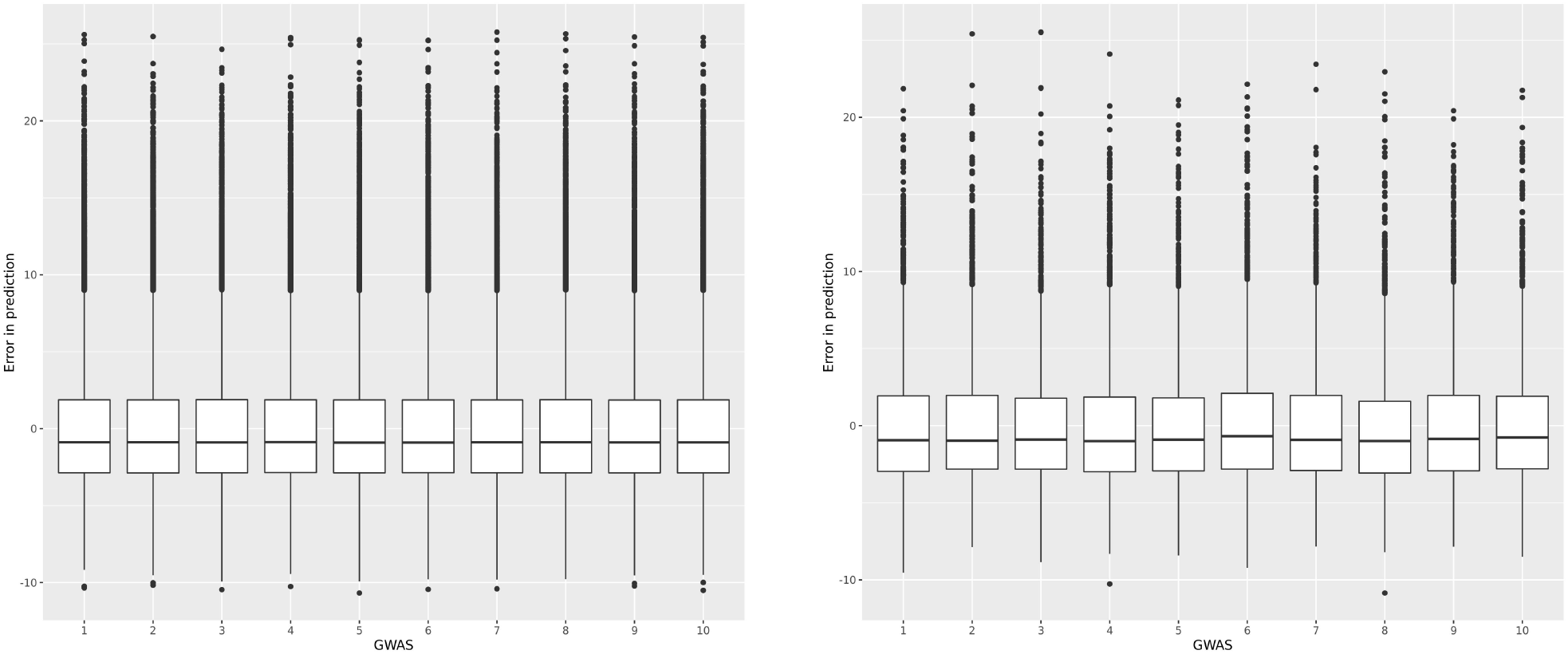
Predicted pre-pregnancy BMI performance on test and training sets. Left panel, the boxplot of the difference between the predicted pre-pregnancy BMI and the observed pre-pregnancy BMI on each training set, using a P-value threshold of 10^−8^. Right panel, the boxplot of the difference between the predicted pre-pregnancy BMI and the observed pre-pregnancy BMI on each test set, using a P-value threshold of 10^−8^.

**Figure 25:**
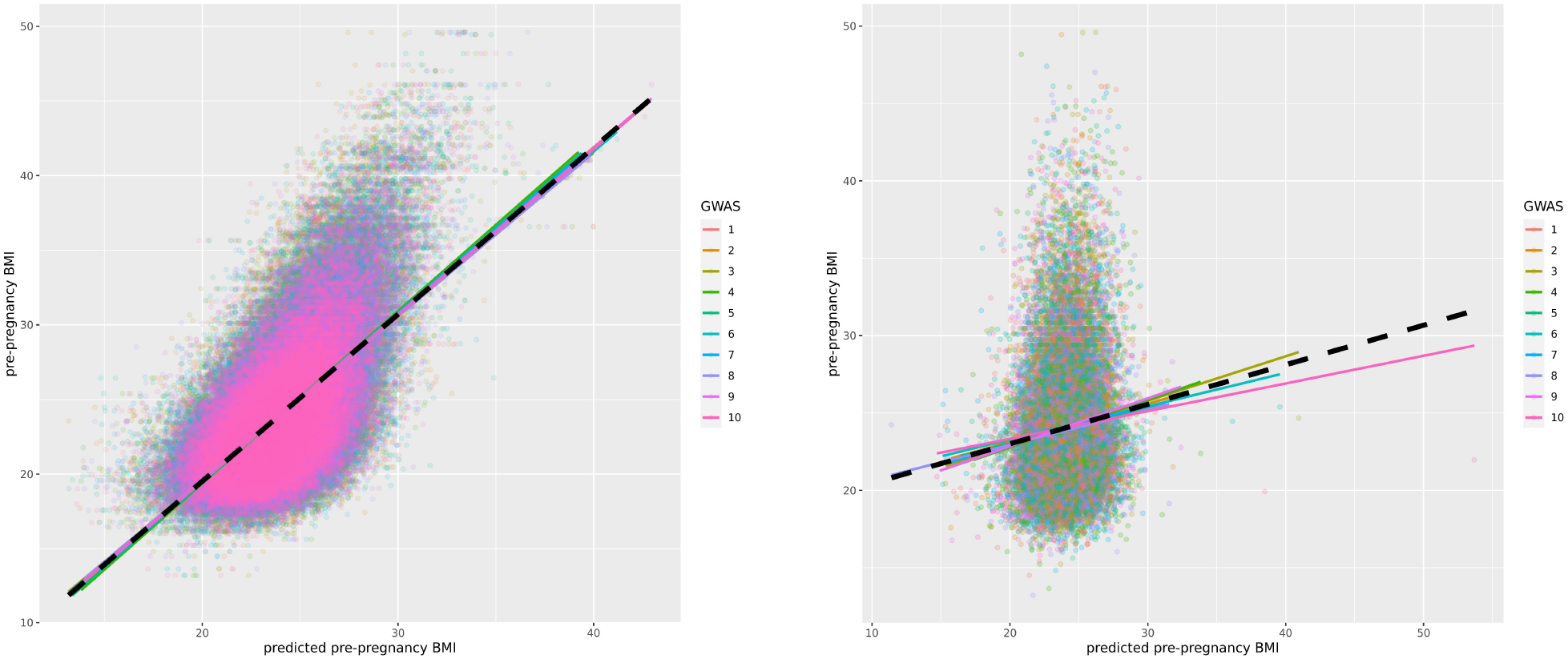
Predicted pre-pregnancy BMI performance on test and training sets. Left panel, the bivariate plot of the predicted pre-pregnancy BMI on training sets against true values using a P-value threshold of 10^−3^. Right panel, the bivariate plot of the predicted pre-pregnancy BMI on test sets against the true values using a P-value threshold of 10^−3^.

**Figure 26:**
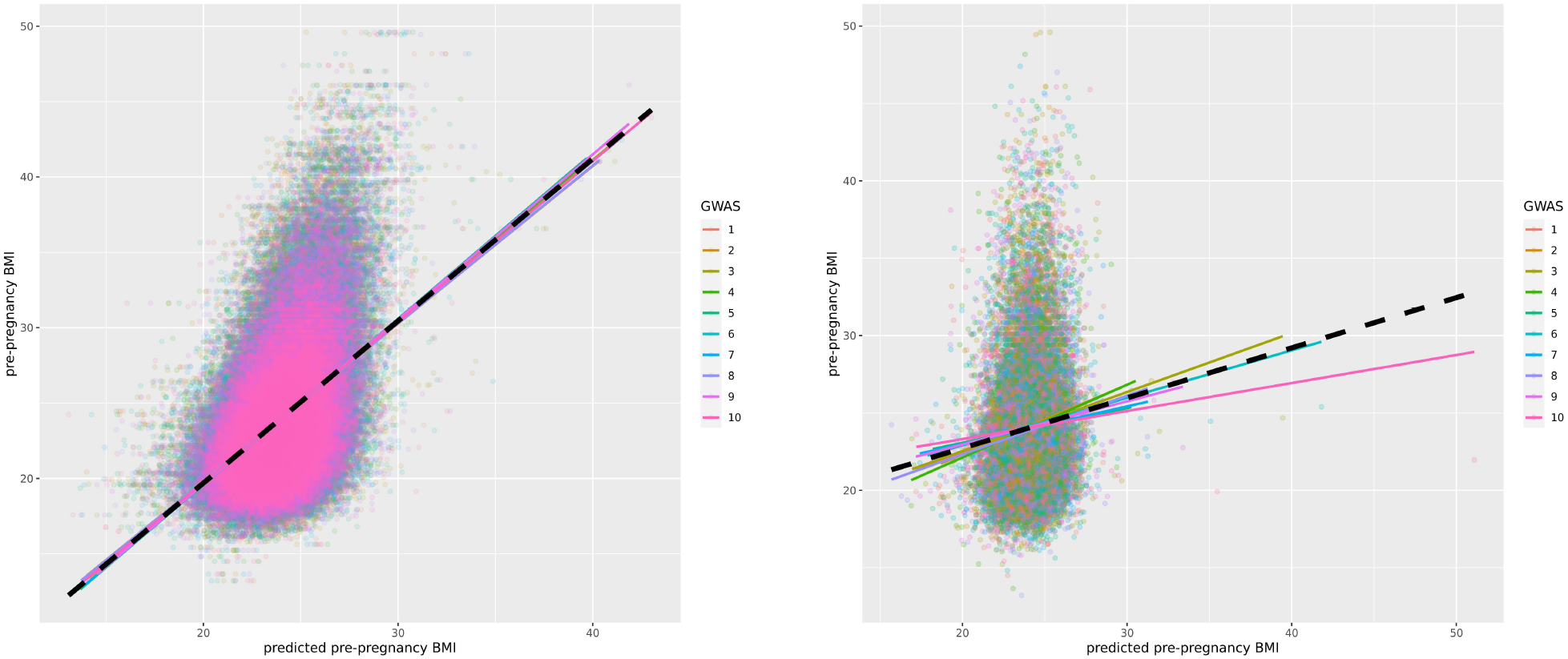
Predicted pre-pregnancy BMI performance on test and training sets. Left panel, the bivariate plot of the predicted pre-pregnancy BMI on training sets against true values using a P-value threshold of 10^−4^. Right panel, the bivariate plot of the predicted pre-pregnancy BMI on test sets against true values using a P-value threshold of 10^−4^.

**Figure 27:**
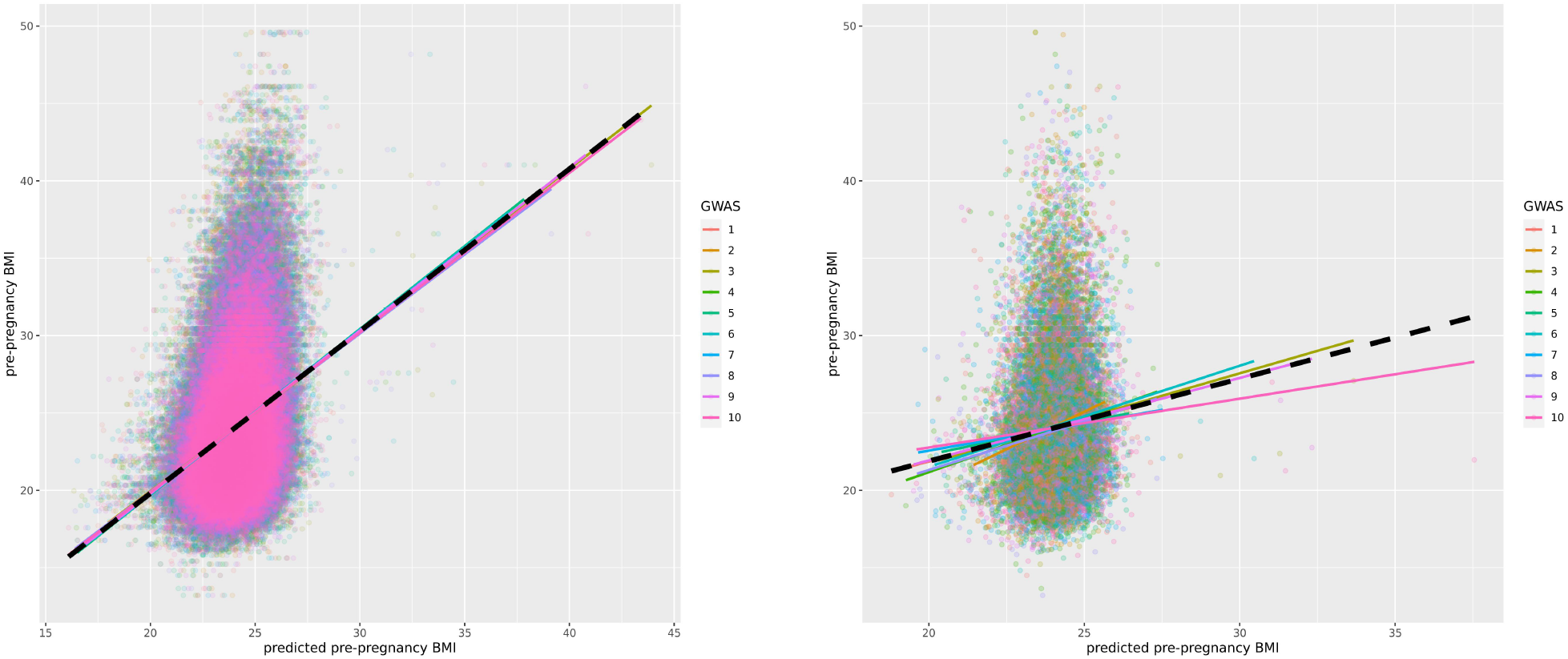
Predicted pre-pregnancy BMI performance on test and training sets. Left panel, the bivariate plot of the predicted pre-pregnancy BMI on training sets against true values using a P-value threshold of 10^−5^. Right panel, the bivariate plot of the predicted pre-pregnancy BMI on test sets against true values using a P-value threshold of 10^−5^.

**Figure 28:**
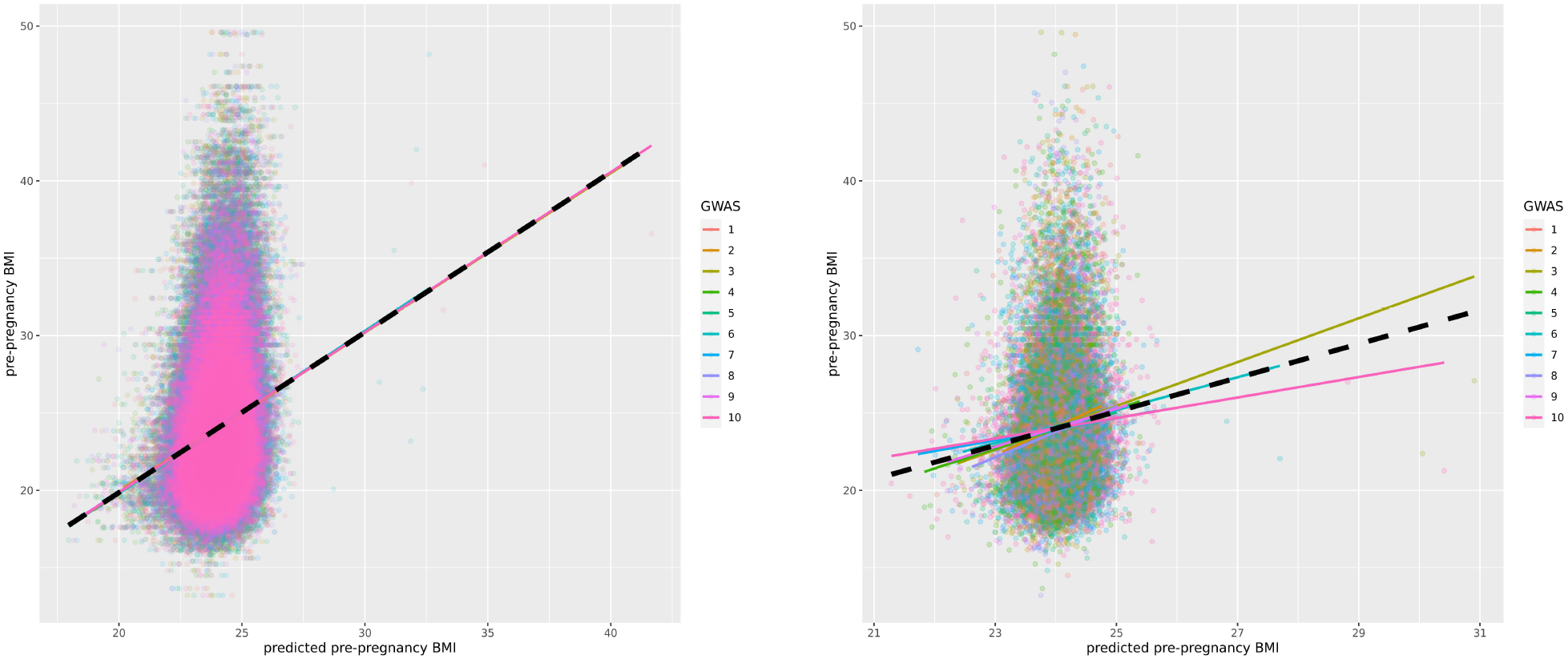
Predicted pre-pregnancy BMI performance on test and training sets. Left panel, the bivariate plot of the predicted pre-pregnancy BMI on training sets against true values using a P-value threshold of 10^−6^. Right panel, the bivariate plot of the predicted pre-pregnancy BMI on test sets against true values using a P-value threshold of 10^−6^.

**Figure 29:**
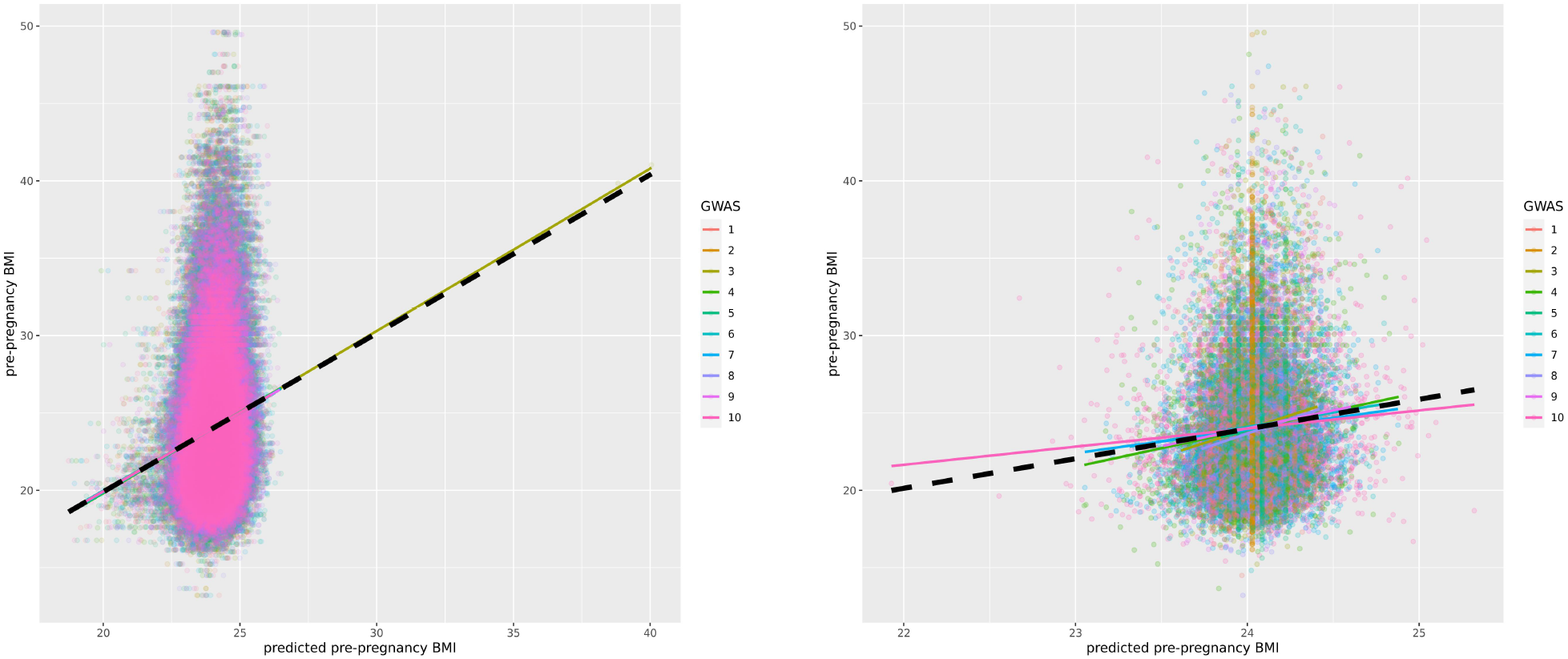
Predicted pre-pregnancy BMI performance on test and training sets. Left panel, the bivariate plot of the predicted pre-pregnancy BMI on training sets against true values using a P-value threshold of 10^−7^. Right panel, the bivariate plot of the predicted pre-pregnancy BMI on test sets against true values using a P-value threshold of 10^−7^.

**Figure 30:**
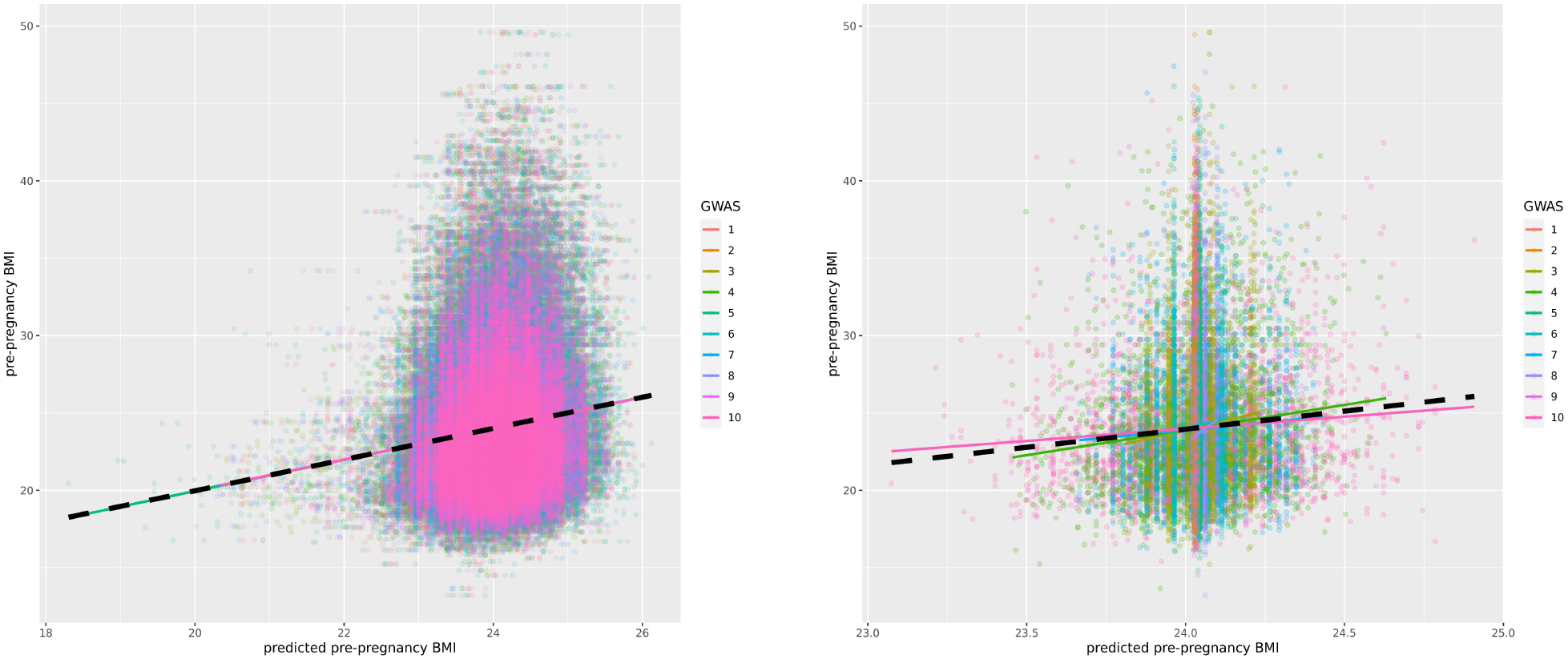
Predicted pre-pregnancy BMI performance on test and training sets. Left panel, the bivariate plot of the predicted pre-pregnancy BMI on training sets against true values using a P-value threshold of 10^−8^. Right panel, the bivariate plot of the predicted pre-pregnancy BMI on test sets against true values using a P-value threshold of 10^−8^.

**Figure 31:**
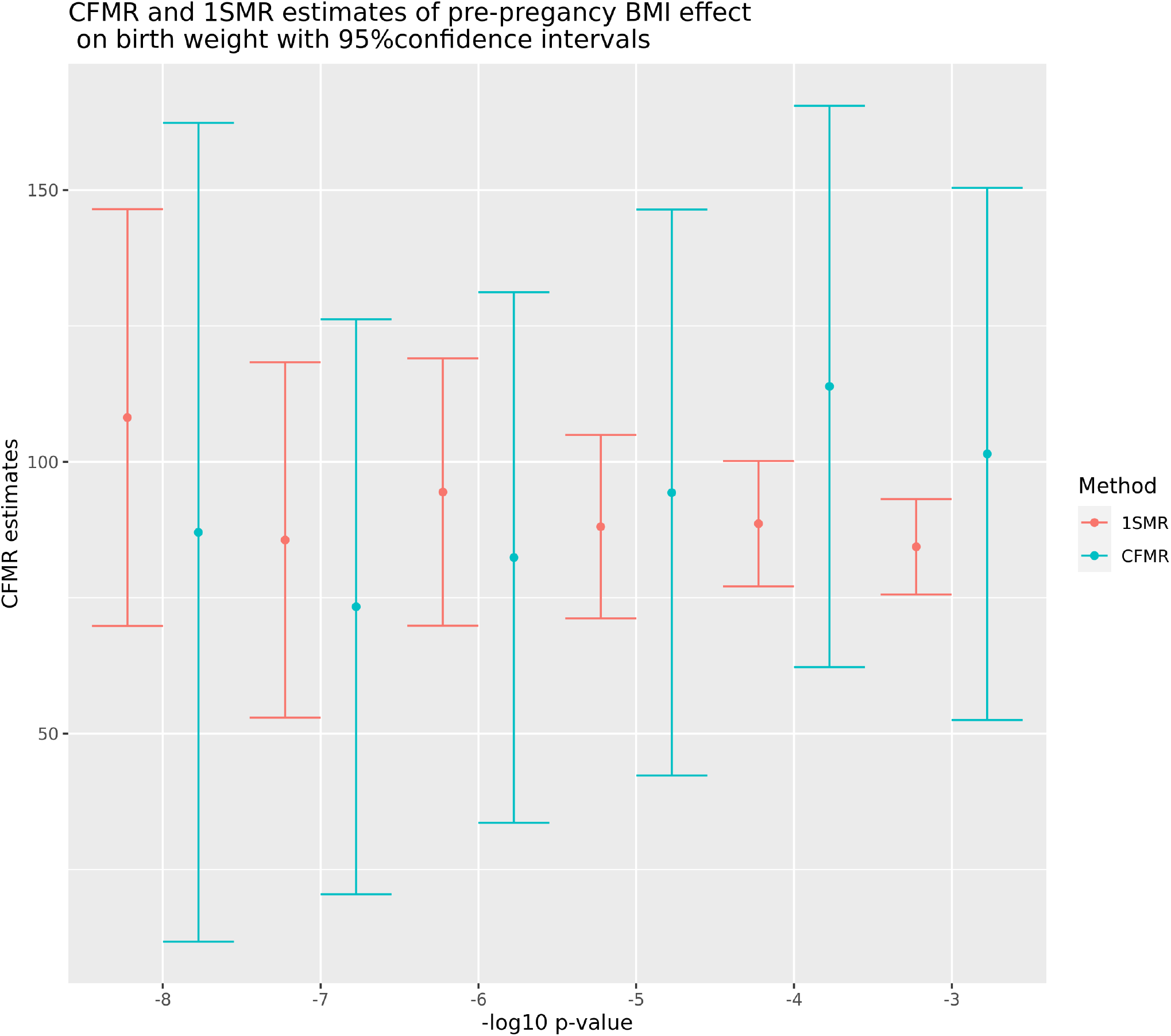
CFMR and one-sample MR (1SMR) estimates of the effect of pre-pregnancy maternal BMI on birth weight, with 95% confidence intervals.

**Table 5:** CFMR estimates (raw scale).

| $-\log_{10}$ SNP<br>P-value | Variance<br>explained by CFI | CFMR<br>estimate | Std. Error<br>error | P-value | 95% CI<br>lower limit | 95% CI<br>upper limit |
| --- | --- | --- | --- | --- | --- | --- |
| -3 | 1.112 % | 24.130 | 5.938 | 0.00005 | 12.491 | 35.768 |
| -4 | 1.102 % | 27.031 | 6.257 | 0.00002 | 14.767 | 39.295 |
| -5 | 1.101 % | 22.399 | 6.308 | 0.00038 | 10.035 | 34.762 |
| -6 | 1.112 % | 19.571 | 5.910 | 0.00093 | 7.986 | 31.155 |
| -7 | 0.951 % | 17.420 | 6.409 | 0.00657 | 4.859 | 29.980 |
| -8 | 0.044 % | 20.669 | 9.125 | 0.02351 | 2.785 | 38.553 |

**Figure 32:**
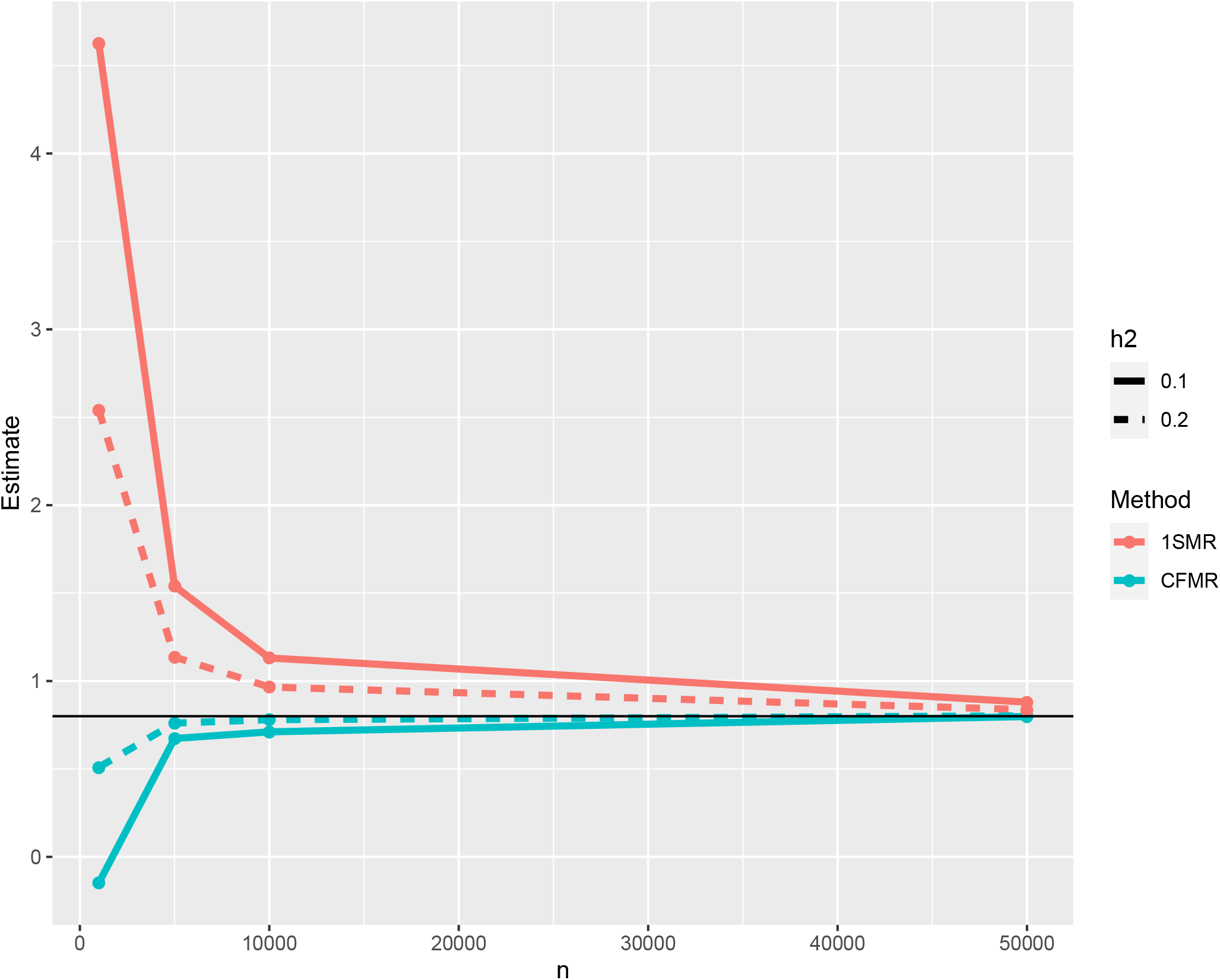
Summary of the simulations performed in Supplementary Section 3.2. The x-axis corresponds to the number individual used in each simulation (1000; 5, 000; 10, 000; and 50, 000), the y-axis corresponds to the estimated effect. The solid horizontal black line corresponds to the true value of the effect to be estimated. The different types of lines correspond to the variance *X* explained by the genetic marker used as instrument (10% and 20%). The red lines (1SMR) correspond to the estimations based on one-sample MR and the green lines correspond to the estimations based on CFMR.

**Table 6:** Association of CFI with potential confounders.

| $-\log_{10}$ SNP P-value | Confounder | Estimate | Std. Error | P-value |
| --- | --- | --- | --- | --- |
| -3 | Mother's age | -0.012 | 0.015 | 0.4254063 |
| -3 | Mother's education | -0.014 | 0.007 | 0.0548912 |
| -3 | Gestational age | 0.017 | 0.005 | 0.0014255 |
| -3 | Mother's smoking | 0.033 | 0.008 | 0.0000443 |
| -4 | Mother's age | -0.004 | 0.020 | 0.8356098 |
| -4 | Mother's education | -0.005 | 0.009 | 0.6290257 |
| -4 | Gestational age | 0.027 | 0.007 | 0.0001246 |
| -4 | Mother's smoking | 0.039 | 0.011 | 0.0002364 |
| -5 | Mother's age | -0.029 | 0.033 | 0.3819641 |
| -5 | Mother's education | 0.008 | 0.016 | 0.6245407 |
| -5 | Gestational Age | 0.051 | 0.012 | 0.0000125 |
| -5 | Mother's smoking | 0.089 | 0.018 | 0.0000005 |
| -6 | Mother's age | -0.087 | 0.066 | 0.1878742 |
| -6 | Mother's education | 0.003 | 0.031 | 0.9251442 |
| -6 | Gestational Age | 0.068 | 0.023 | 0.0034211 |
| -6 | Mother's smoking | 0.121 | 0.035 | 0.0005351 |
| -7 | Mother's age | -0.105 | 0.131 | 0.4222781 |
| -7 | Mother's education | 0.027 | 0.061 | 0.6548937 |
| -7 | Gestational Age | 0.075 | 0.046 | 0.1035003 |
| -7 | Mother's smoking | 0.003 | 0.069 | 0.9634737 |
| -8 | Mother's age | -0.372 | 0.227 | 0.1006788 |
| -8 | Mother's education | -0.052 | 0.106 | 0.6247780 |
| -8 | Gestational age | 0.088 | 0.079 | 0.2663506 |
| -8 | Mother's smoking | -0.001 | 0.120 | 0.9932419 |

**Table 7:** Estimation of the effect of X on Y for *β* = 0.8 by one sample MR and CFMR, respectively. The simulations are detailed in Supplementary Section 3.2.

| N | $h^2$ | CFMR $\hat{\beta}$<br>estimate | 1SMR $\hat{\beta}$<br>estimate |
| --- | --- | --- | --- |
| 1000 | 0.10 | -0.15 | 4.63 |
| 5000 | 0.10 | 0.67 | 1.54 |
| 10000 | 0.10 | 0.71 | 1.13 |
| 50000 | 0.10 | 0.80 | 0.88 |
| 1000 | 0.20 | 0.51 | 2.54 |
| 5000 | 0.20 | 0.76 | 1.14 |
| 10000 | 0.20 | 0.78 | 0.97 |
| 50000 | 0.20 | 0.80 | 0.84 |

